# Synergistic Modulation of Macrophages by Methotrexate and RELA siRNA Folate-Liposome: A Precision Therapy to Prevent Joint Degradation in Collagen-Induced Arthritic Rats

**DOI:** 10.1101/2024.01.03.574006

**Authors:** Simran Nasra, Dhiraj Bhatia, Ashutosh Kumar

**Author notes:** Corresponding Author: Ashutosh Kumar, Associate Professor Biological & Life Sciences, School of Arts & Sciences, Ahmedabad University, Central Campus, Navrangpura, Ahmedabad 380009 Gujarat, India.

## Abstract

Rheumatoid arthritis (RA) is a chronic autoimmune disorder characterized by inflammation and joint destruction. Current treatments, such as Methotrexate (MTX), while effective, often have therapeutic limitations like high plasma C_max_ and lack of sustained release. This study explores a synergistic approach to RA therapy using folate-liposomal co-delivery of MTX and RELA siRNA, aimed at RAW264.7 macrophage repolarization through inhibition of the NF-κB pathway. Extensive invitro characterizations demonstrate the stability and biocompatibility of this combinatorial therapy in folate-liposomes. In collagen-induced arthritis (CIA) rat model, we observed a reduction in synovial inflammation and improved mobility following treatment. The combined MTX and RELA siRNA approach indirectly inhibits inflammatory cytokines and other biochemical parameters such as Rheumatoid factor (RF) and C-reactive protein (CRP). The targeted macrophage delivery yields a marked therapeutic effect in RAW264.7 murine macrophages, potentially modulating the M1 to M2 macrophage polarization. Overall, this research presents a promising avenue for innovative therapies in RA management by inhibiting the inflammatory cascade and preventing joint damage.

## 1. Introduction

Rheumatoid arthritis (RA) is a pervasive autoimmune disorder characterized by chronic inflammation that leads to joint destruction, causing debilitating pain, disability, and compromised quality of life. [1] Its global prevalence remains substantial, with approximately 23 million of the world’s population grappling with this challenging condition. [2] As a leading cause of disability worldwide, RA exerts a significant burden on patients and their families, healthcare systems, and economies. [3] It is imperative to continue exploring innovative therapeutic strategies that can enhance the quality of life for those living with RA and alleviate the societal and economic impact of the disease. [4] The management of RA has evolved significantly over the years, with Methotrexate (MTX) serving as a fundamental cornerstone in its treatment [5]. MTX, an immunosuppressive drug, has been a pivotal tool in dampening the immune response responsible for joint inflammation and destruction [6]. However, despite its effectiveness, MTX has various therapeutic limitations. Some patients experience inadequate relief, intolerable side effects, or the development of resistance over time [7]. The side effects are found to be correlated with dose escalation due to its low bioavailability and short half-life [8]. Furthermore, the complexity of RA’s pathogenesis extends beyond the confines of a single treatment [9]. As such, researchers have continued to investigate novel and multifaceted approaches that can provide more comprehensive relief to those afflicted by RA [10]. Among the therapeutic avenues that have garnered significant attention, the role of macrophages in RA pathogenesis has come to the forefront [11]. Macrophages, as innate immune cells, exhibit remarkable plasticity and heterogeneity in response to local microenvironmental cues [12]. Classically activated M1 macrophages are instrumental in initiating and sustaining inflammation, playing a central role in the tissue damage observed in RA [13]. Conversely, alternatively, activated M2 macrophages are tasked with resolving inflammation and promoting tissue repair [14]. Understanding the role of these macrophage phenotypes and their plasticity has opened new possibilities for therapeutic intervention in RA [15].

This study presents a promising approach to RA therapy by combining MTX with RELA siRNA, a small interfering RNA targeting the NF-κB, encapsulated in folate-liposomes. Folate receptor-targeted liposomes (FOL-liposomes), offer a means of precise drug delivery, ensuring that the therapeutic agents reach their intended destination [16]. In the case of this study, macrophages, central players in the RA inflammatory cascade, are the primary target. These FOL-liposomes have undergone rigorous characterization, encompassing evaluations of size, morphology, stability, and biocompatibility, to ensure their suitability as a drug delivery system. This comprehensive assessment provides a robust foundation for their therapeutic use. Employing a collagen-induced arthritis (CIA) rat model, the results offer promising prospects, as the study observes a significant reduction in synovial inflammation, coupled with marked improvements in the mobility of the affected rats. This innovative approach indirectly inhibits vital inflammatory cytokines, such as tumor necrosis factor (TNF), interleukin-6 (IL-6), and interleukin-1 beta (IL-1β), while also reducing levels of rheumatoid factor (RF) and C-reactive protein (CRP).

The distinctive feature of this strategy lies in its ability to repolarize macrophages, promoting the transition from a proinflammatory M1 phenotype to a pro-resolving M2 phenotype, observed in RAW264.7 cells. This shift offers a unique therapeutic advantage, as M2 macrophages are known for their anti-inflammatory properties, contributing to tissue repair and homeostasis. By influencing this phenotypic change, the therapy holds the potential to curb inflammation and prevent further joint damage. The potential to shift macrophage phenotypes, alleviate inflammation, and prevent joint damage represents a significant step forward in the quest to improve the lives of those affected by this challenging autoimmune disorder.

## 2. Results and discussion

### 2.1 Optimization of the MTX + RELA siRNA FOL-liposomes formulation using 3-factor and 3-level Box–Behnken response surface design (3^3^BBD)

Optimizing the parameters involved in the preparation of liposomal formulations is essential for enhancing the liposomes’ physicochemical properties and their ability to effectively transport therapeutic payloads to their designated location of activity [17]. The Box-Behnken design is a type of response surface methodology (RSM) used in experimental design to study the relationship between multiple independent variables and their impact on multiple responses [18]. It is particularly useful when the relationships between variables and responses are nonlinear and difficult to predict. In this context, a three-factor, three-level Box-Behnken response surface design (3^3^BBD) was employed for formulation optimization. The responses were individually fitted, including entrapment efficiency (%EE), particle size, and ζ-potential of MTX + RELA siRNA FOL-liposomes. The objective was to identify the most appropriate model with the highest adjusted and prediction R-squared values (adjusted R-squared values ≥ 0.9). Model terms that did not show statistical significance were removed to achieve a higher R-squared (R²) value. The Design Expert software generated a design matrix based on data collected from 15 experimental runs. As indicated in Table 1, the quadratic model was chosen for all three responses. An analysis of variance (ANOVA) was conducted with a significance level of p < 0.05. Following the simplification of the model, the ultimate equations describing the factors and interactions for the responses were generated using Design-Expert Version 12.0.1.0 (Stat-Ease Inc., Suite 480, Minneapolis, MN, USA) in terms of coded variables, as shown in Table 1 and Supplementary Table S1.

**Table 1.** Three independent variables (A,B, and C) and its coded values fitted in Design-Expert.

| Optimization of MTX liposomes |  | Actual amount at coded values |  |  |
| --- | --- | --- | --- | --- |
| Coded values |  | -1 | 0 | +1 |
| Variables | (A) DSPC/CHOL | 2:1 | 4:1 | 6:1 |
|  | (B) Lipid/MTX | 5 | 10 | 15 |
|  | (C) DSPE-PEG-FOL | 0.02 | 0.04 | 0.06 |

#### 2.1.1 Effect of the independent variables on Entrapment Efficiency (%EE)

The equation provided below represents a second-order polynomial response surface model with three independent variables, denoted as A, B, and C. The equation is used to predict the response variable %EE based on different levels of the independent variables. Each term in the equation represents a specific factor or interaction effect:

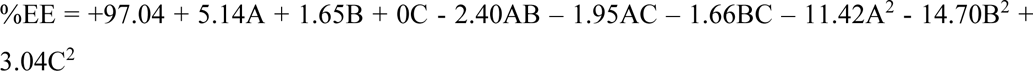

Where: %EE is the predicted response variable. A, B, and C are the independent variables (factors) representing different experimental conditions or levels. The coefficients associated with each term represent the main effects and interaction effects of the independent variables. Factor A, representing the DSPC/CHOL ratio, exerts a positive linear effect on %EE, signifying that an increase in this ratio enhances %EE. However, its negative quadratic term suggests that beyond an optimal point, further increases may decrease %EE. Factor B, the Lipid/MTX ratio, also exhibits a positive linear relationship with %EE, indicating that a higher lipid-to-MTX ratio improves %EE, while its negative quadratic term implies an optimal point for this ratio. The interaction terms, −2.40AB and −1.95AC, illustrate that the presence of other factors can modify the influence of Factor A. Notably, Factor C (DSPE-PEG-FOL concentration) does not exhibit a linear impact on %EE, as indicated by its coefficient of 0, yet its positive quadratic term (3.04C^2^) suggests the presence of an optimal concentration for enhancing %EE (shown in Figure 1).

**Figure 1.**
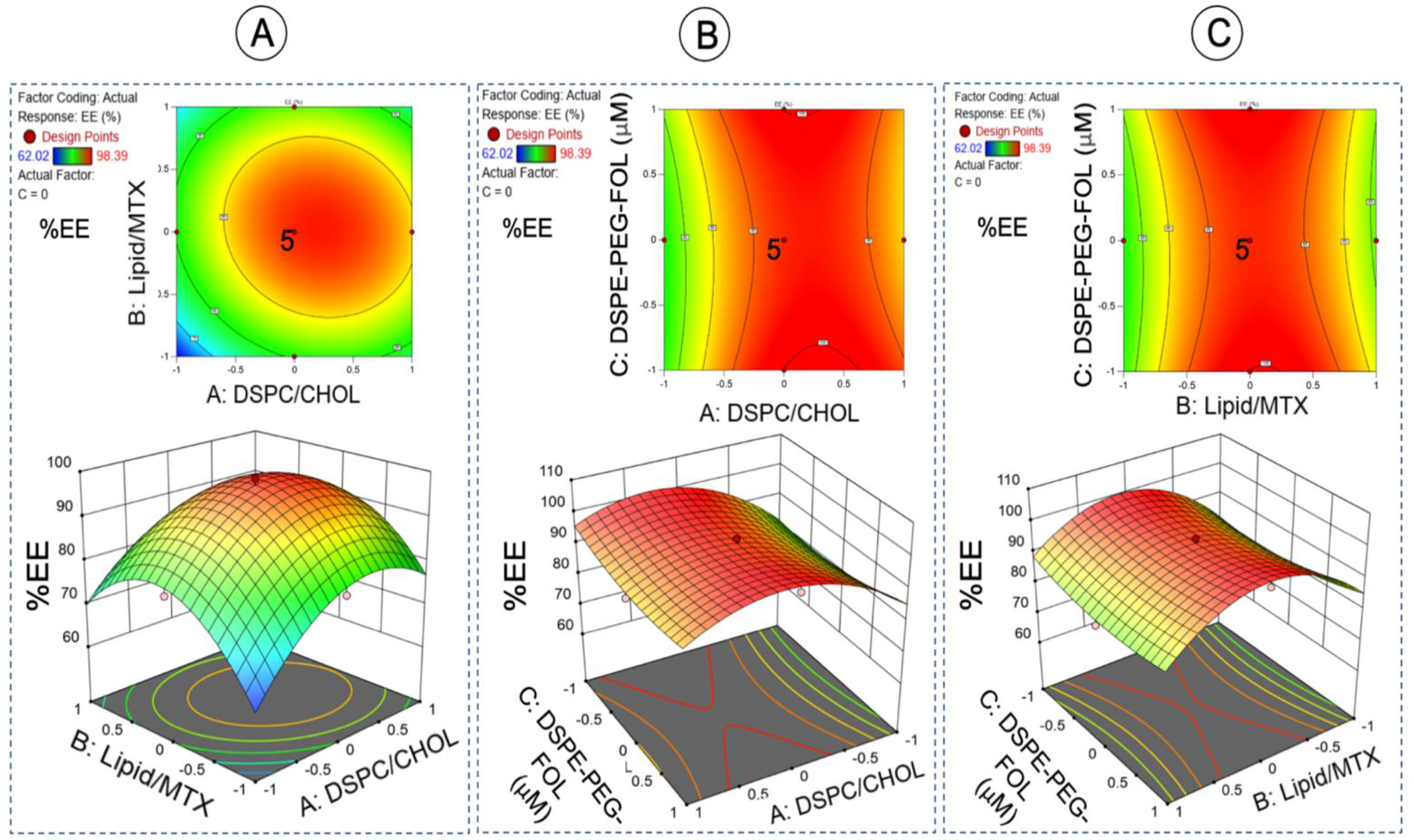
Two and Three-dimensional surface plots for the main effects and interactions of the independent factors of MTX + RELA siRNA FOL-liposomes formulae (A: DSPC/CHOL molar ratio, B: Lipid/MTX molar ratio, and C: DSPE-PEG-FOL concentration) on EE%.

#### 2.1.2 Effect of the independent variables on Particle Size

Particle size can directly affect the delivery of drugs, where the size of particles can influence bioavailability, tissue targeting, and therapeutic outcomes. The equation for Particle Size offers a comprehensive understanding of how the factors DSPC/CHOL (Factor A), Lipid/MTX (Factor B), and DSPE-PEG-FOL concentration (Factor C) influence particle size in the formulation [19].

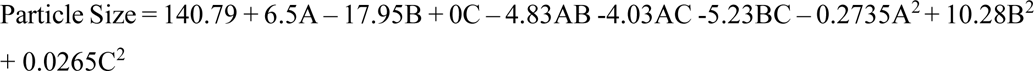

Factor A, representing the DSPC/CHOL ratio, has a positive linear effect, indicating that higher levels of DSPC/CHOL lead to larger particle sizes. However, its negative quadratic term suggests an optimal ratio beyond which size reduction may occur. Factor B, representing the Lipid/MTX ratio, exhibits a negative linear effect, implying that increasing this ratio results in smaller particle sizes. Interaction terms (AB, AC, and BC) demonstrate that the presence of one factor can modulate the impact of another on particle size (as shown in Figure 2). Achieving the desired particle size entails finding the right balance between the linear and quadratic effects of Factors A, B, and C to optimize formulation conditions effectively. The coefficients for the quadratic terms (A^2^, B^2^, C^2^) represent the curvature or nonlinearity of the response surface to the respective variables. For example, the negative coefficient for A^2^ (- 0.2735) suggests that increasing A may have a diminishing return on the Particle Size response, while the positive coefficient for B^2^ (10.28) indicates that increasing B leads to an increase in the Particle Size response.

**Figure 2.**
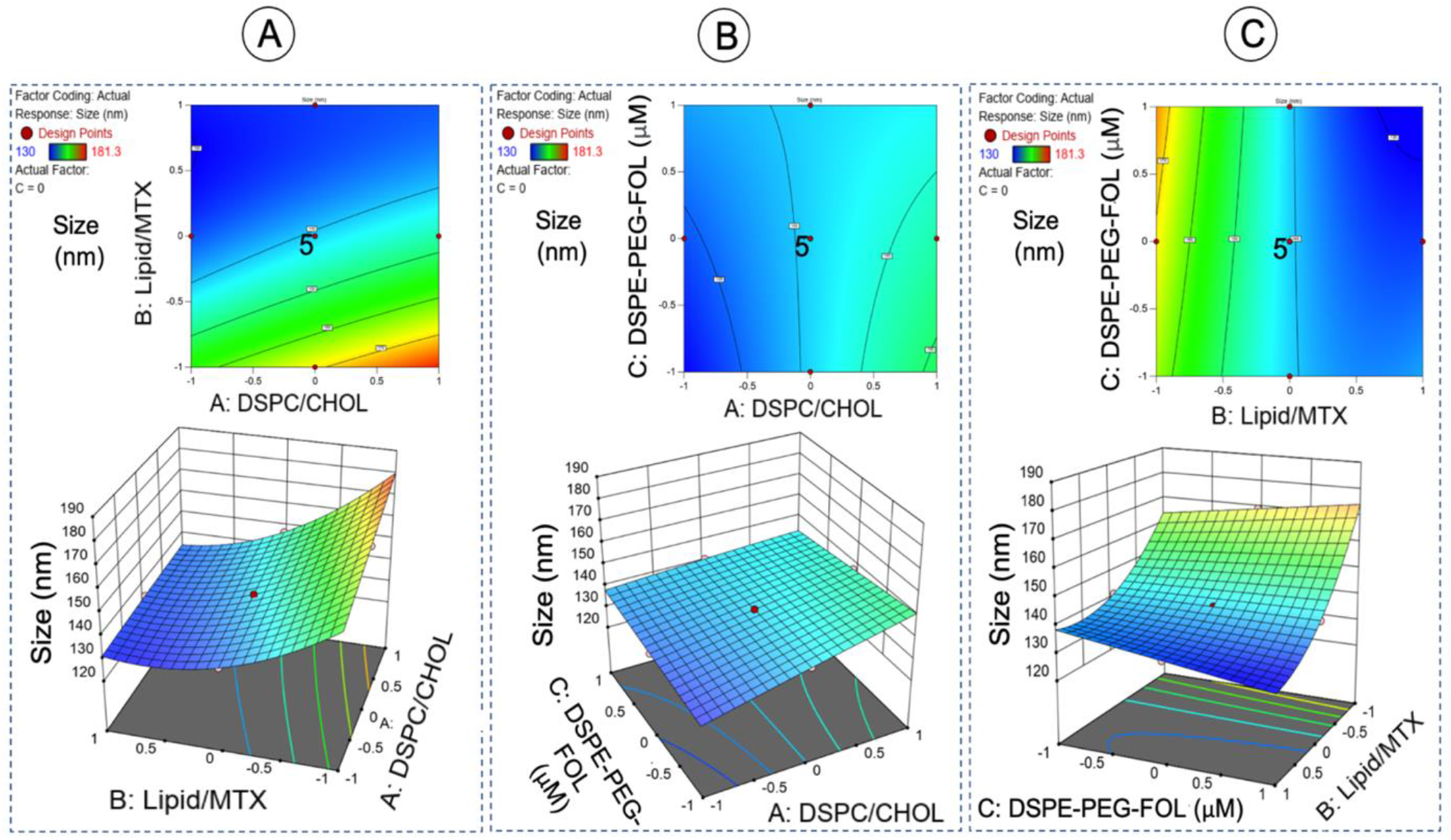
Two and Three-dimensional surface plots for the main effects and interactions of the independent factors of MTX + RELA siRNA FOL-liposomes formulae (A: DSPC/CHOL molar ratio, B: Lipid/MTX molar ratio, and C: DSPE-PEG-FOL concentration) on Particle Size.

**Figure 3.**
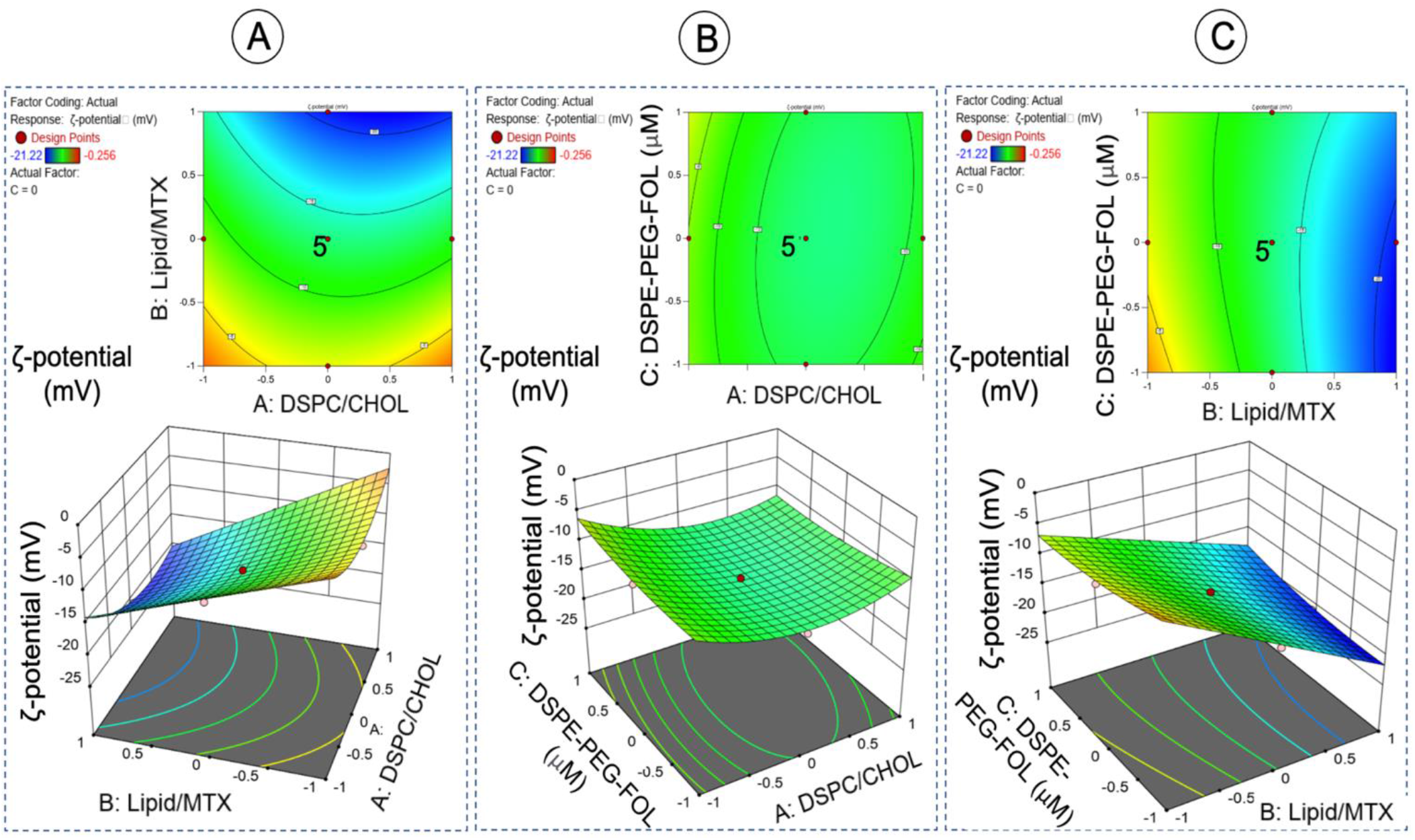
Two and Three-dimensional surface plots for the main effects and interactions of the independent factors of MTX + RELA siRNA FOL-liposomes formulae (A: DSPC/CHOL molar ratio, B: Lipid/MTX molar ratio, and C: DSPE-PEG-FOL concentration) on ζ-potential.

#### 2.1.2 Effect of the independent variables on ζ-potential

Understanding how these factors affect ζ-potential is particularly relevant in fields such as colloid chemistry and nanoparticle engineering, where surface charge plays a pivotal role in colloidal stability, electrostatic interactions, and applications like drug delivery.

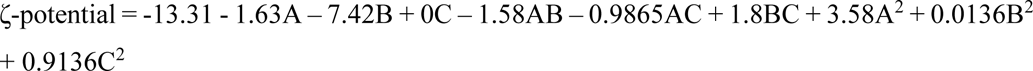

Factor A, reflecting the DSPC/CHOL ratio, demonstrates a negative linear effect, indicating that higher levels of DSPC/CHOL contribute to a lower ζ-potential. The presence of a negative quadratic term for A implies an optimal ratio where ζ-potential reaches a minimum. In contrast, Factor B, representing the Lipid/MTX ratio, has a negative linear effect, signifying that higher Lipid/MTX ratios lead to a reduced ζ-potential. The positive quadratic term for B suggests an optimal ratio for minimizing surface charge. Factor C exhibits no linear effect on ζ-potential, but the presence of a positive quadratic term indicates an optimal concentration that positively influences the surface charge. The coefficients for the quadratic terms (A^2^, B^2^, C^2^) represent the curvature or nonlinearity of the response surface with respect to the respective variables. The interaction terms (AB, AC, and BC) reveal that the combined presence of factors modulates the effect on ζ-potential.

The best-predicted ratio for DSPC/CHOL is 4:1, 10 for Lipid/MTX, and 0.06 for DSPE-PEG-FOL (as depicted in Supplementary Figure S1). Formulation 5, exhibits 98.39 % EE, 140.8 nm Hydrodynamic size, and −12.9 ζ-potential. The overall desirability value (D-Overall) ranges from 0 to 1, with higher values indicating that the experimental conditions are closer to achieving all desired targets for the responses and independent variables. An overall desirability value of 1 implies that the experimental conditions are optimal for all responses. The most desirable formulation was with DSPC/CHOL = 4:1, Total Lipids/MTX = 10, and DSPE-PEG-FOL = 0.06.

### 2.2 Unveiling the Characteristic Features of Formulated MTX + RELA siRNA FOL-Liposomes

Evaluating a successful Methotrexate (MTX) encapsulation within liposomal formulations is a pivotal step in the drug delivery system. As shown in Figures 4 A and B, MTX exhibited a distinct absorbance peak at 303 nm, which served as a reliable indicator of its presence within the liposomes. Significantly, a dose-dependent increase in absorbance was observed, allowing for the construction of a precise calibration curve. This curve facilitated the quantification of MTX levels within the liposomes, vital for dose control and assessment of the therapeutic efficacy. Notably, empty liposomes did not manifest the characteristic 303 nm absorbance peak, further substantiating the specific encapsulation of MTX. Both bare MTX and MTX- loaded liposomes exhibited this peak, providing unequivocal evidence of successful encapsulation. Additionally, the formation of the final combinatorial liposome system was meticulously validated through advanced imaging techniques, namely Transmission Electron Microscopy (TEM) and Dynamic Light Scattering (DLS).

**Figure 4.**
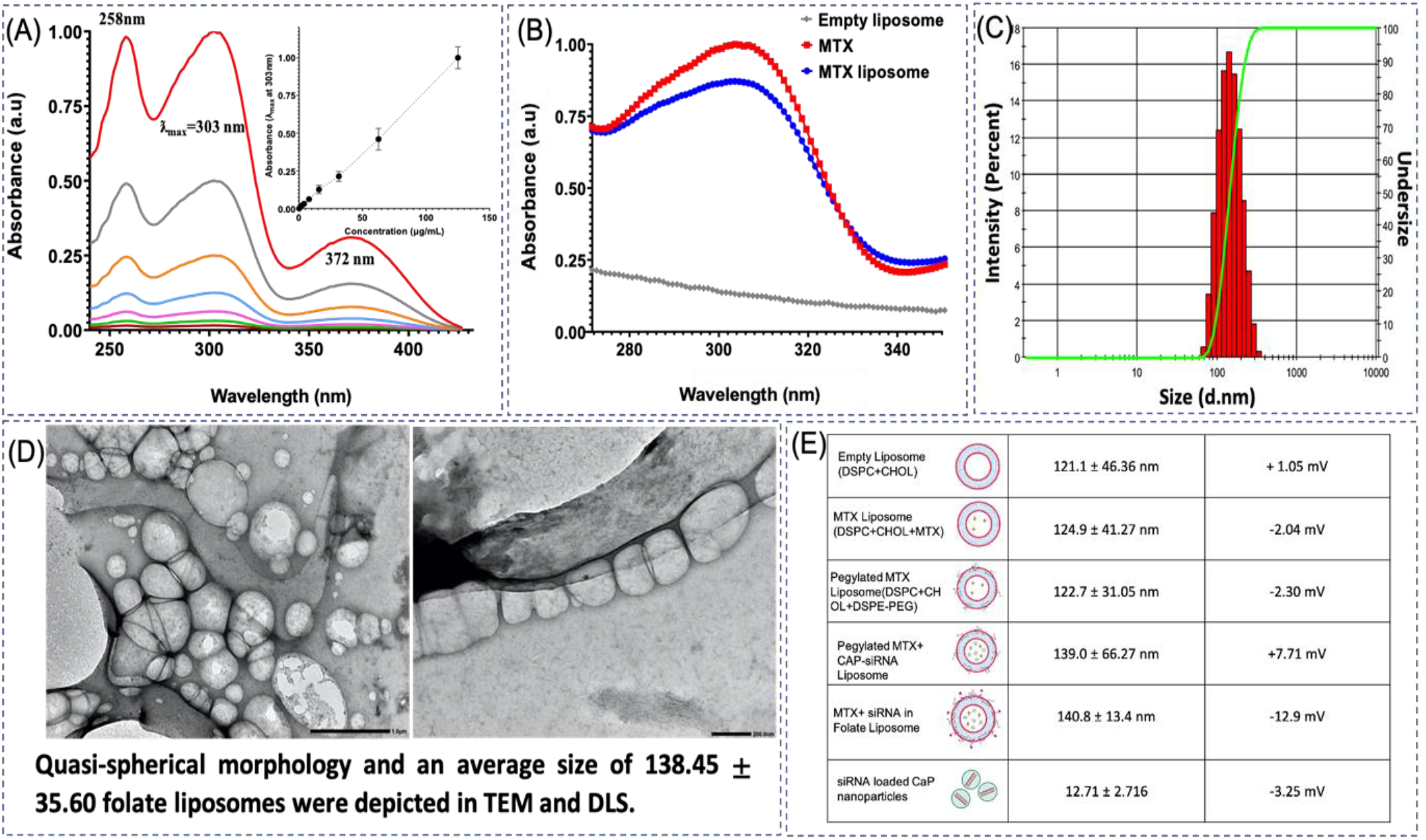
Representative images for (A) Absorbance maxima observed at 303 nm for MTX measured at different concentrations and its standard curve is plotted, (B) Successful encapsulation of MTX in liposomes based on its absorbance maxima, (C) Hydrodynamic diameter of 140.8 ± 66.27 nm for MTX + RELA siRNA FOL-liposomes, (D) TEM images show uniform and quasi-spherical morphology of MTX + RELA siRNA FOL-liposomes before and after downsizing, and (E) Hydrodynamic size and zeta potential of all the individual and combinatorial nanoparticles.

Remarkably, the final formulation displayed a noticeable increase in hydrodynamic size in comparison to empty liposomes, a feature with substantial implications for drug release kinetics and therapeutic performance, as shown in Figure 4C. Moreover, the more negative zeta potential of the final formulation indicated improved stability, attributed to electrostatic repulsion among the constituent particles. This attribute is pivotal for maintaining the integrity and homogeneity of the liposomal formulation over time, especially in biological environments. Lastly, TEM analysis revealed a quasi-spherical morphology and an average size of 138.45 ± 35.60 nm for folate liposomes, which are pertinent characteristics in terms of biocompatibility and interaction with target cells (as shown in Figure 4D). This comprehensive analysis underscores the successful development and characterization of the combinatorial liposome systems (Shown in Figure 4E), reaffirming its suitability for targeted drug delivery.

As observed in the AFM analysis (Figure 5 A and B), the functionalization of folate onto the surface of liposomes can be visually confirmed. Upon careful analysis of these images, distinctive features or additional protrusions can be observed on the liposome surface, indicating the presence of folate molecules. The functionalized liposomes, with folate molecules attached, appear to exhibit characteristic changes compared to regular liposomes. The AFM images may show small, distinct irregularities distributed across the liposome surface, which correspond to the folate molecules. Folate receptors are often overexpressed on the surface of inflammatory cells. By functionalizing liposomes with folate, they can specifically target and bind to these cells, enhancing drug delivery and therapeutic efficacy.

**Figure 5.**
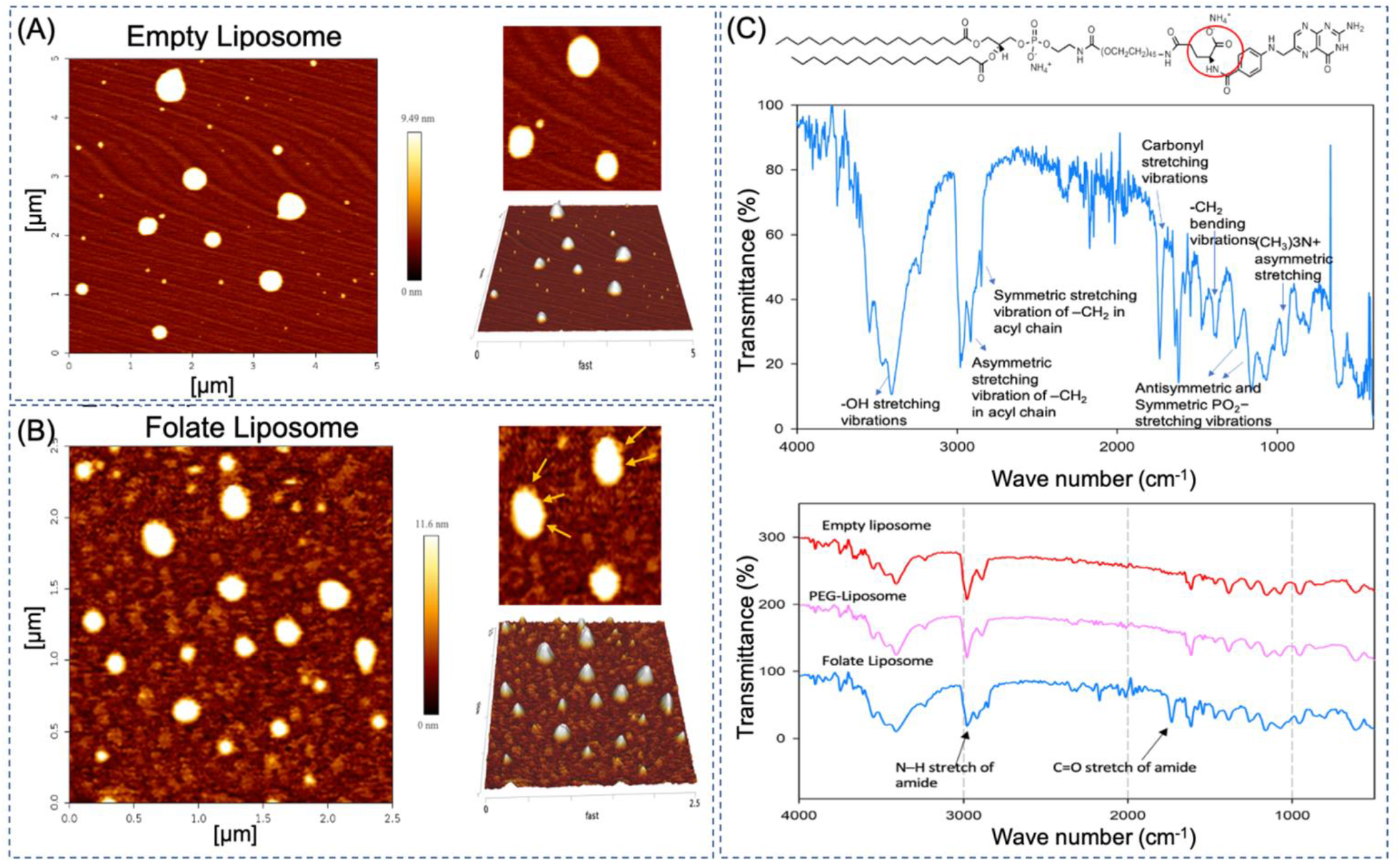
Representative images for (A) Nanoliposomes without folate conjugation, (B) Nanoliposomes with folate conjugation (FOL-liposome), yellow arrows indicate the surface irregularities and folate conjugation on liposome surface (C) FTIR peaks obtained in FOL-liposome, (D) Peculiar confirmatory peaks of DSPE-PEG-FOL obtained in FOL-liposome, suggesting successful functionalization of folate on the surface of liposomes.

Additionally, as depicted in Figure 5 C, the addition of CO and NH peaks in Fourier transform infrared (FTIR) spectra occurs due to the formation of an amide bond resulting from the conjugation of folate molecules with DSPE-PEG (1,2-distearoyl-sn-glycero-3-phosphoethanolamine-N-[amino(polyethylene glycol)]). This reaction involves the formation of an amide bond (CO-NH) between these functional groups, resulting in the attachment of folate to the liposome surface. The presence of the amide bond is detected using FTIR spectroscopy. In an FTIR spectrum, different functional groups in a sample absorb infrared radiation at characteristic frequencies, leading to the appearance of peaks in the spectrum. The -CO and -NH peaks in the FTIR spectrum of the functionalized liposomes are indicative of the newly formed amide bond. The CO peak usually appears in the region between 1630 cm-1 and 1820 cm-1, and the NH peak typically appears around 3300 cm-1 [20]. These peaks are strong indicators of the successful conjugation of folate to DSPE-PEG, confirming the presence of the amide bond.

The addition of PEG (polyethylene glycol) to liposomes can enhance their stability, leading to maintained size and zeta potential over an extended period, such as 30 days (as shown in Figure 6 A and B). As a result, the liposomes are less likely to aggregate or be recognized by the immune system, leading to improved stability over time. The utilization of calcium phosphate nanoparticles as carriers for siRNA, subsequently loaded into liposomes, provides a stable and protective environment for siRNA, shielding it from enzymatic degradation and premature release. The calcium phosphate core acts as a reservoir for siRNA, facilitating controlled and sustained release. Importantly, calcium phosphate is a material naturally present in the human body, reducing concerns about toxicity or immune responses. As shown in Figure 6 C, MTX liposomes show a lack of distinct peaks in the XRD pattern, which was previously observed in bare MTX, suggesting an amorphous state. The amorphous nature of methotrexate in liposomes can be beneficial for drug delivery purposes, as amorphous formulations often offer improved solubility, dissolution, and bioavailability compared to crystalline forms. The liposome encapsulation of methotrexate can enhance its stability and allow for controlled release, while the amorphous state promotes better drug release kinetics.

**Figure 6.**
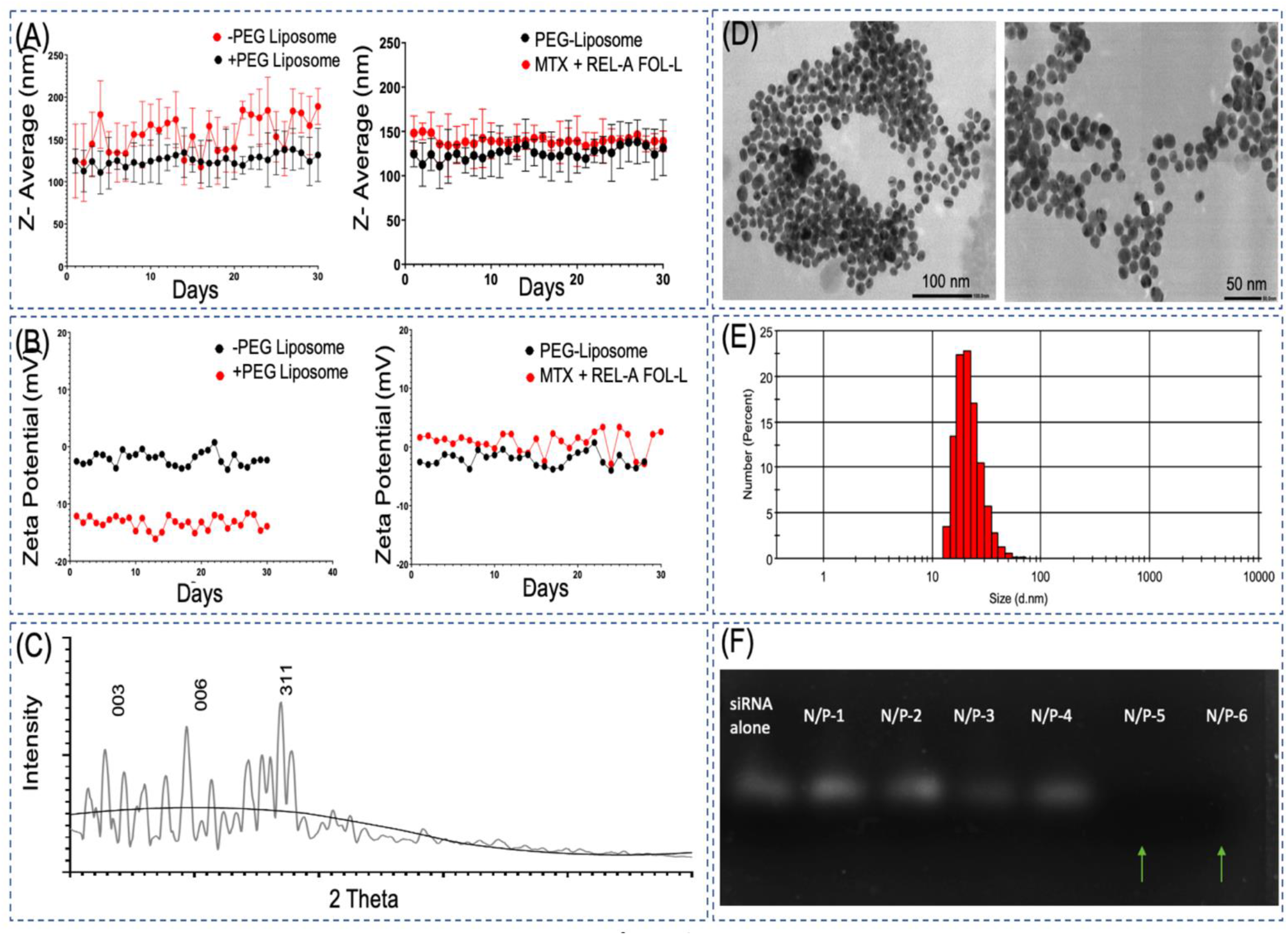
Graphical representation for (A, B) Stability of liposomes after PEGylation and addition of folate on the surface, depicted by not much variation in their hydrodynamic size and zeta potential, (C) Methotrexate liposomes (black line) show a lack of distinct peaks in the XRD pattern, which was previously observed in MTX (grey peaks), suggesting an amorphous or disordered state, (D, E) Morphological analysis in TEM and hydrodynamic size for Calcium Phosphate nanoparticles and (F) the N/P ratio obtained after 4.

As observed in Figure 6 D and E, calcium phosphate nanoparticles in the range of 10-50 nm using TEM indicates that these nanoparticles possess small dimensions and fall within a relatively uniform size range. Additionally as validated in Figure 6 F, an N/P ratio of 5 is used in the final formulation. The N/P ratio refers to the ratio of the number of nitrogen (N) atoms in the cationic component of a nanoparticle formulation to the number of phosphate (P) groups in the nucleic acid cargo, such as siRNA or plasmid DNA. This ratio is crucial for the efficient encapsulation and delivery of nucleic acids in nanoparticle-based gene delivery systems. By using an N/P ratio of 5, we have optimized the formulation to achieve a balance between effective encapsulation and delivery of the nucleic acids, in this case, likely to be siRNA targeting NF-kB, and minimizing potential toxicity or instability.

### 2.3 Drug Entrapment and Kinetic Release Study

As depicted in Figure 7 (upper panel), the retention time of MTX was observed at 1.942 minutes in the HPLC chromatogram, confirming the presence of MTX in the liposomes. The entrapment efficiency reflects the percentage of the total MTX used in the formulation that is successfully encapsulated within the liposomes. In this case, it was found that 98.39% of the total MTX used was effectively encapsulated in the liposomes. High entrapment efficiency is desired in drug delivery systems as it ensures a higher proportion of the drug is protected and delivered to the target site, leading to improved therapeutic efficacy.

**Figure 7.**
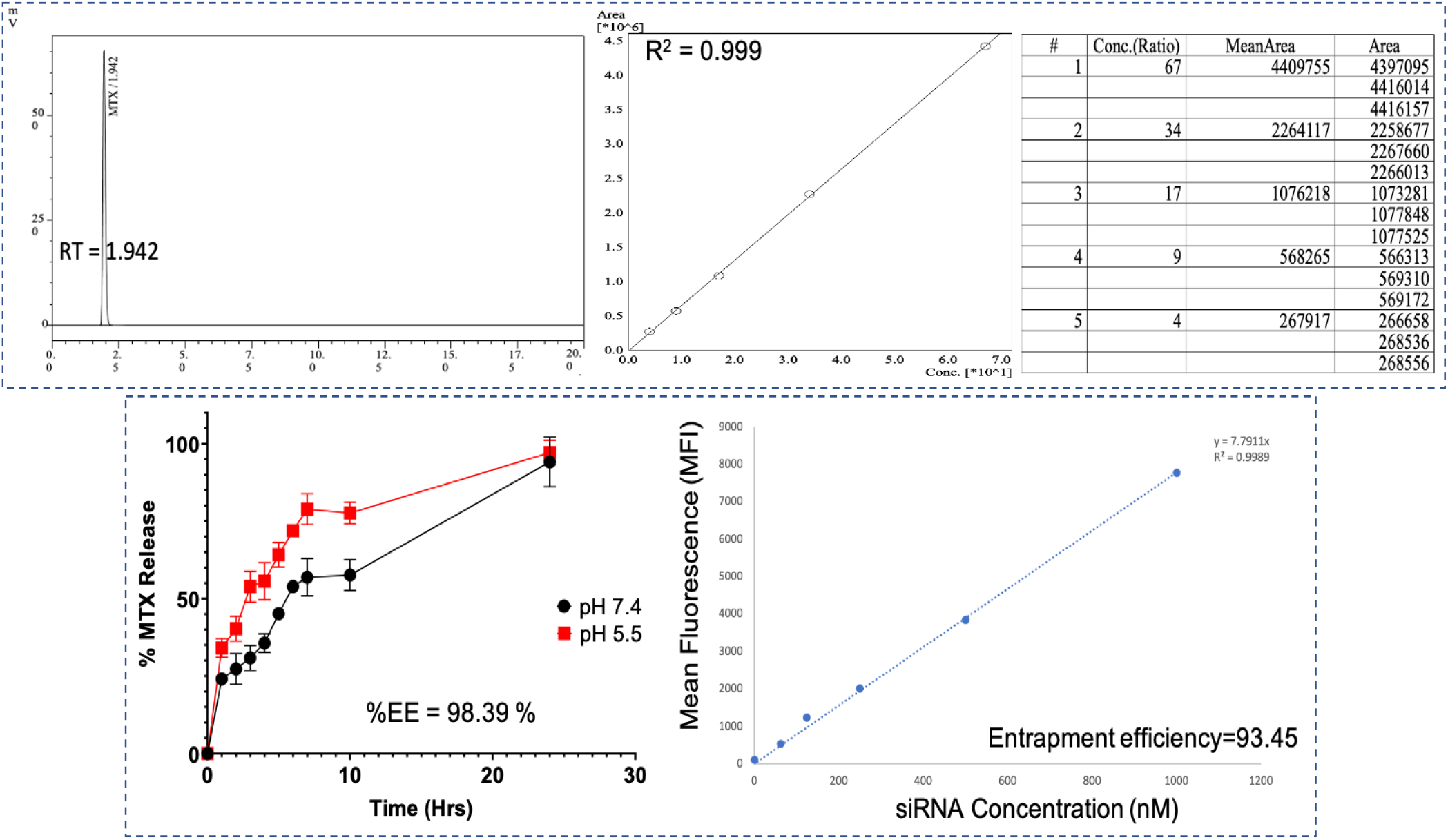
The upper panel suggests the retention time peak for MTX is at 1.942 minutes, obtained in HPLC, its calibration curve, and standards used in this analysis. The lower panel suggests % entrapment efficiency and release kinetics of MTX at pH 7.4 and pH 5.5, and %EE of RELA siRNA obtained at 93.45 % using known siRNA standards.

The observation that MTX had a more efficient and sustained release at acidic pH compared to pH 7.4 suggests that the liposomes are designed for targeted drug delivery (as shown in Figure 7, lower panel). Under acidic conditions, such as those found in intracellular compartments, the liposomes may undergo drug release more readily, resulting in improved drug delivery to the target site. This selective release in macrophages enhances therapeutic efficacy while reducing potential side effects on healthy tissues. The entrapment efficiency of RELA siRNA was found to be 93.45% by calculating the mean fluorescence index (MFI), indicating that a significant portion of the siRNA was successfully encapsulated within the liposomes. This high entrapment efficiency ensures effective delivery of the siRNA, allowing for targeted gene silencing and modulation of the RELA gene.

### 2.4 Enhancing Drug Efficacy: Dose Reduction of MTX and REL-A siRNA in MTX + REL-A FOL-Liposome in RAW264.7 Macrophages

The IC_50_ for MTX shifted from 18.03 µM to 1.33 µM when added with REL-A siRNA in FOL-Liposome (shown in Figure A). Also, liposomes facilitate the uptake of siRNA, which alone did not show any effect on cells (Figure 8 B). Dose reduction was observed in combination with MTX and REL-A siRNA FOL-Liposome (shown in Figures 8 C and D). Combining both therapies reduced their individual doses significantly (shown in Figure 8 E). This shift indicates that the combination of MTX with REL-A siRNA in FOL-Liposomes made MTX significantly more potent in inhibiting cell growth. This suggests a potential synergistic effect between MTX and REL-A siRNA, resulting in improved therapeutic outcomes at lower concentrations of MTX.

**Figure 8.**
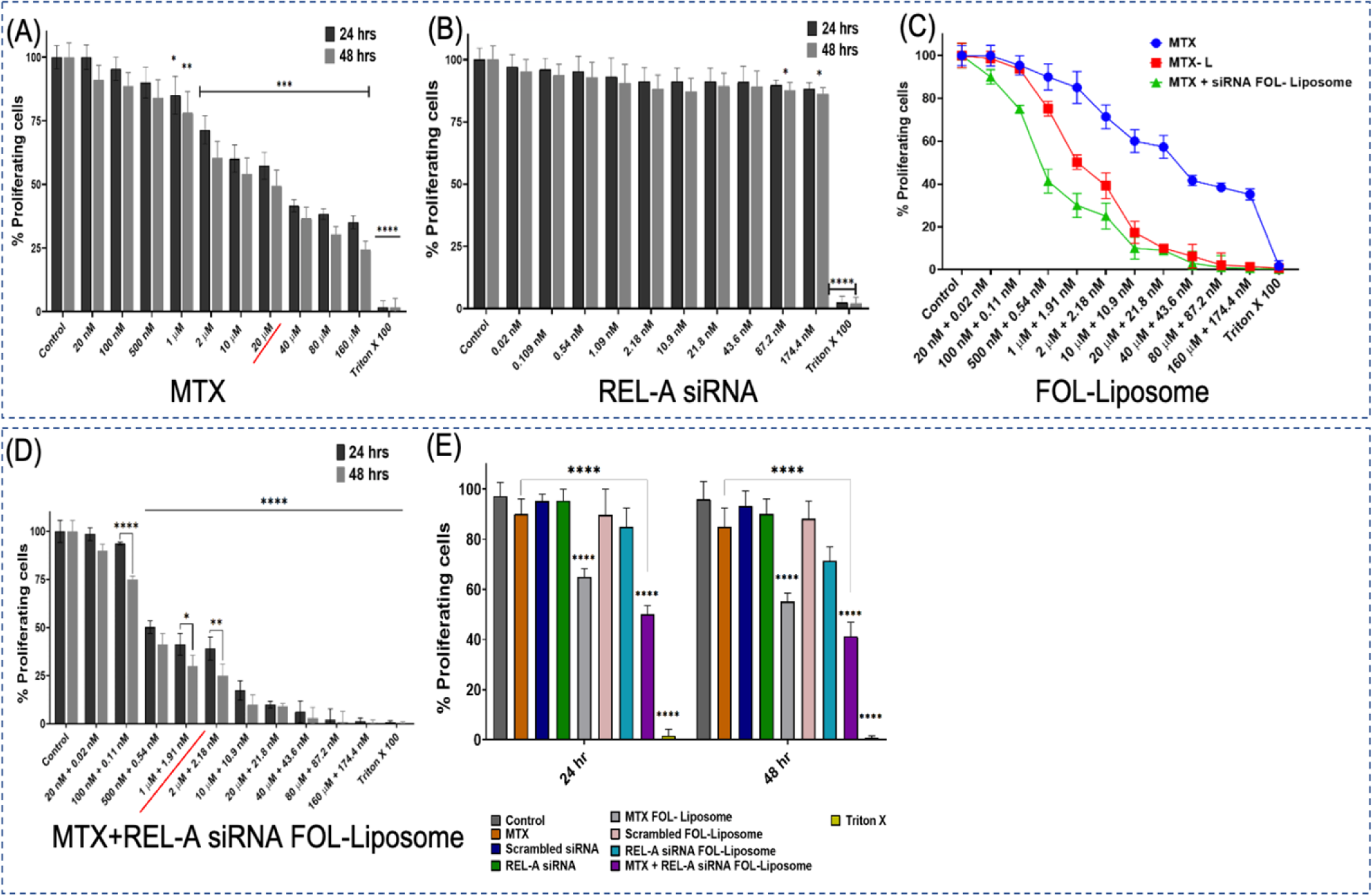
Dose reduction was observed in combination with MTX and REL-A siRNA FOL-Liposome. The IC_50_ for MTX shifted from 18.03 µM to 1.33 µM when added with REL-A siRNA in FOL-Liposome. Combining both therapies reduced their individual doses significantly. Also, liposomes facilitate the uptake of siRNA, which alone did not show any effect on cells. Mean ± SD, * p < 0.05, ** p < 0.01, *** p < 0.001, **** p < 0.0001, respectively.

In the Compusyn studies (depicted in Figure 9), the aim was to investigate the interaction between Methotrexate (MTX) and REL-A siRNA in a liposomal formulation on activated RAW264.7 cells. To determine if the combination of these two agents resulted in a synergistic, additive, or antagonistic effect, they used the combination index (CI) method. The combination index (CI) is a quantitative measure used to assess the combined effect of two drugs. It is calculated based on the dose-response curves of the individual drugs and their combination. The CI value provides information about the nature of the drug interaction: Synergism (CI < 1): A CI value less than 1 indicates synergism, which means that the combined effect of the two drugs is greater than the expected additive effect. Additive Effect (CI = 1): A CI value equal to 1 indicates an additive effect, where the combined effect of the two drugs is exactly as expected by adding their individual effects. There is no synergy or antagonism observed in this case. Antagonism (CI > 1): A CI value greater than 1 indicates antagonism, where the combined effect of the two drugs is weaker than the expected additive effect. This means that the drugs may interfere with each other’s actions, leading to reduced efficacy. In this study, all the treatments showed a CI value less than 1 (CI < 1). This indicates that the combination of MTX and REL-A siRNA in a liposome resulted in a synergistic effect on activated RAW264.7 cells.

**Figure 9.**
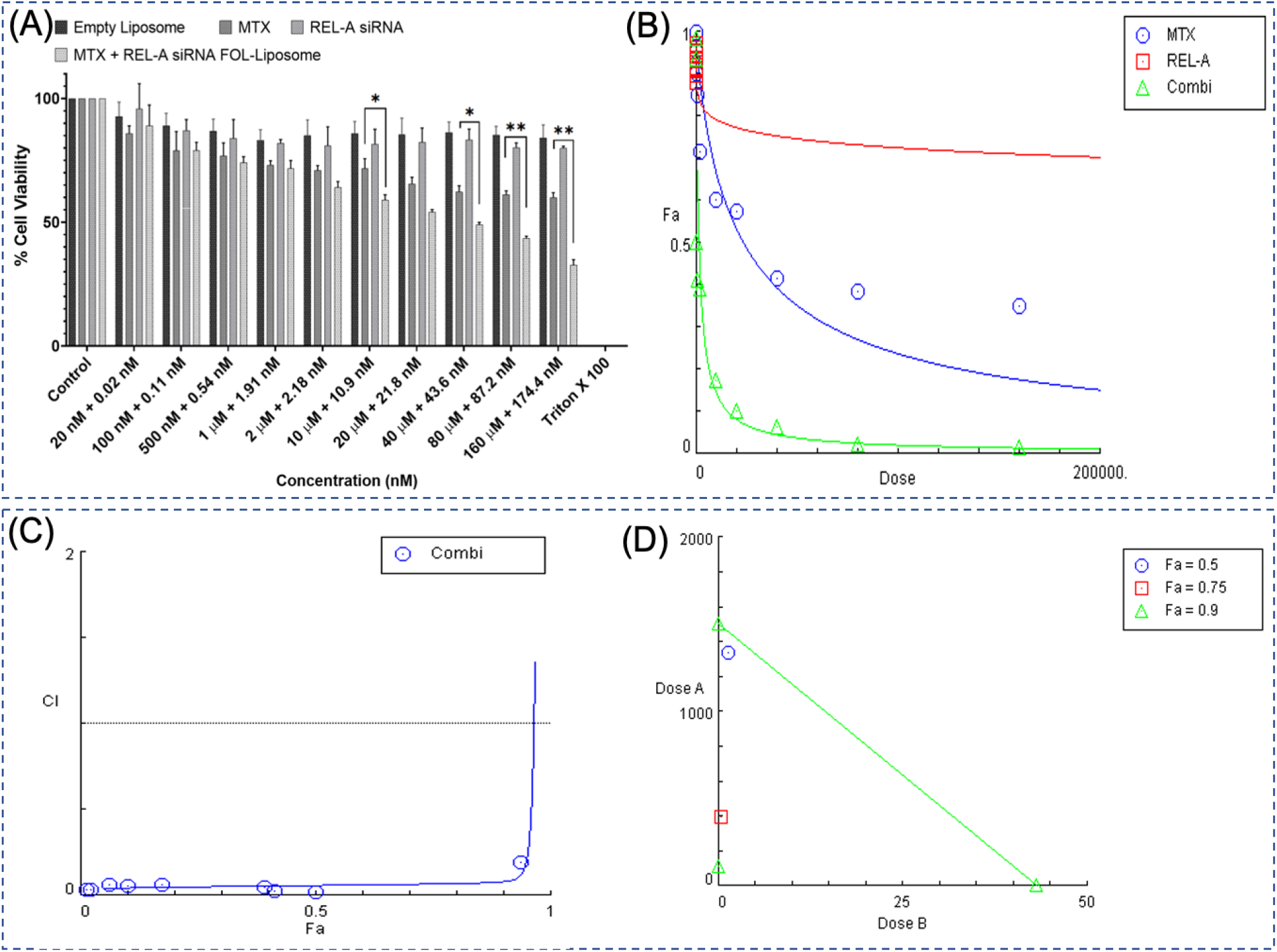
Graphical representation of cell viability assay using Trypan Blue exclusion (A) and Dose-response curve (B), Combination Index (CI) (C), and Fraction Affected (Fa) (D). Fa represents the fraction of cells or organisms affected by a treatment at a given dose. The CI is a quantitative measure that assesses the interaction between multiple drugs or treatments. A CI less than 1, coupled with a Fa indicating a substantial effect at lower doses, would suggest a synergistic interaction, potentially allowing for reduced doses of both MTX and REL-A siRNA while maintaining or even enhancing their therapeutic impact. Mean ± SD, * p < 0.05, ** p < 0.01, *** p < 0.001, **** p < 0.0001, respectively.

### 2.5 Internalization of MTX + RELA siRNA FOL-liposome in RAW264.7 murine macrophages

Figure 10 provides evidence that activated macrophages (M1) show a higher uptake of Rhodamine folate liposomes compared to resting macrophages (M0). The increased uptake is attributed to the overexpression of folate receptors on the surface of activated macrophages. In the context of the figure’s experiment, when these Rhodamine folate liposomes are exposed to both resting and activated macrophages, the activated macrophages, being in the M1 state, exhibit significantly higher uptake of the liposomes compared to the resting macrophages in the M0 state. The specific targeting and higher uptake of Rhodamine folate liposomes by activated macrophages highlight the potential of this approach for targeted drug delivery or therapeutic interventions. By harnessing the folate receptor overexpression in M1 macrophages, it is possible to deliver drugs or therapeutic payloads specifically to the sites of inflammation or diseased tissues, leading to more effective treatments with reduced side effects on healthy cells.

**Figure 10.**
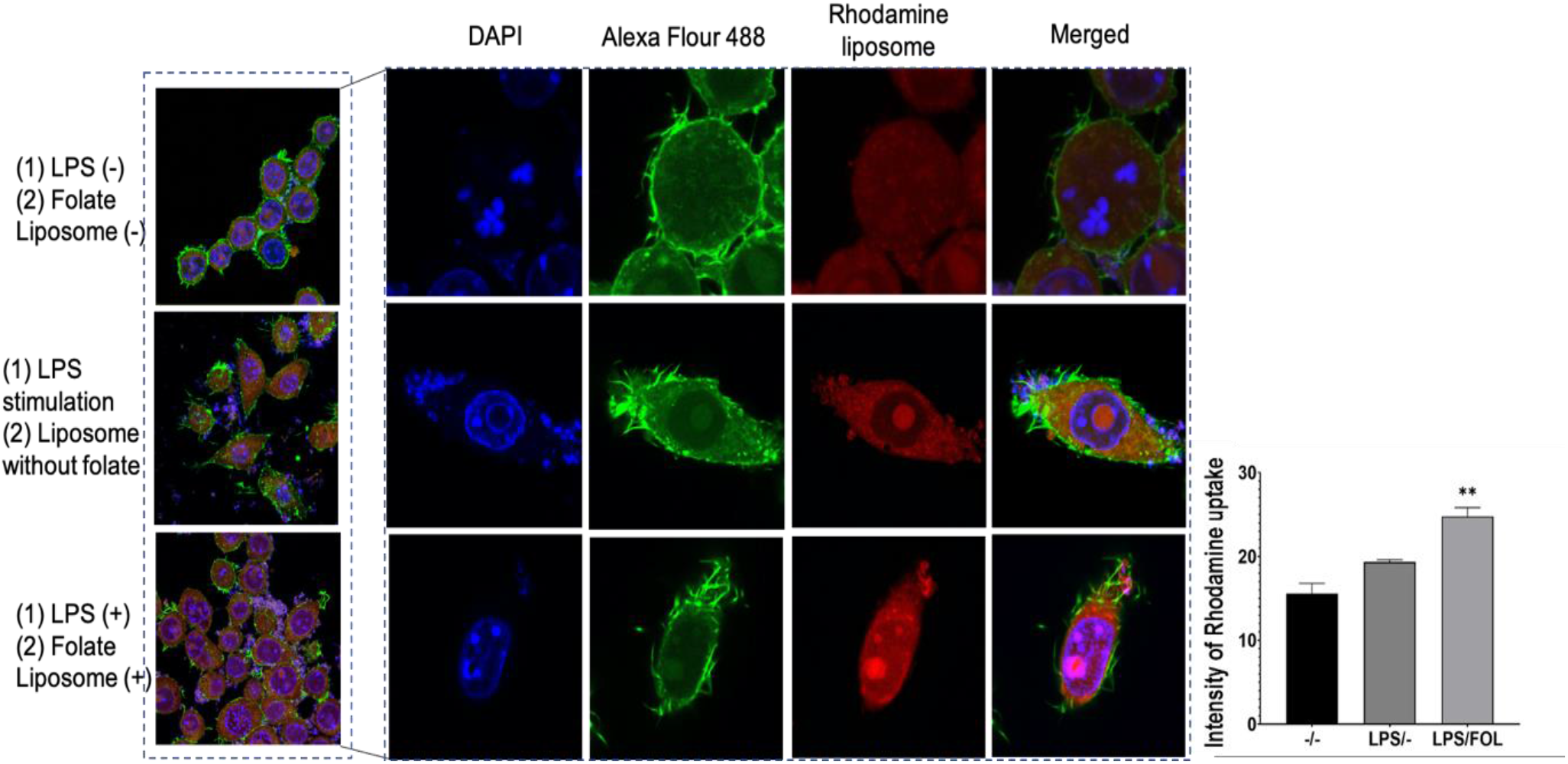
Pictorial and graphical representation of enhanced internalization of MTX + RELA siRNA due to Folate functionalization in naïve macrophages (M0) and activated M1 phenotype. Alexa Flour 488 is used as a cytoplasmic stain and DAPI is used as a nuclear stain, to localize the accumulation of Rhodamine after liposome internalization. Mean ± SD, * p < 0.05, ** p < 0.01, *** p < 0.001, **** p < 0.0001, respectively.

### 2.6 MTX + RELA siRNA induced M1 to M2 macrophage polarization observed in RAW264.7

The induction of M1-polarized RAW264.7 macrophages, characterized by pro-inflammatory and immune-stimulating functions, involves treating the commonly used RAW264.7 macrophage cell line with an inflammatory stimulus, such as LPS. The confirmation of M0 to M1 macrophage switching is evidenced by the detection of CD80 and CD86 positive populations, key co-stimulatory molecules upregulated during M1 activation (Figure 11 A to D).

**Figure 11.**
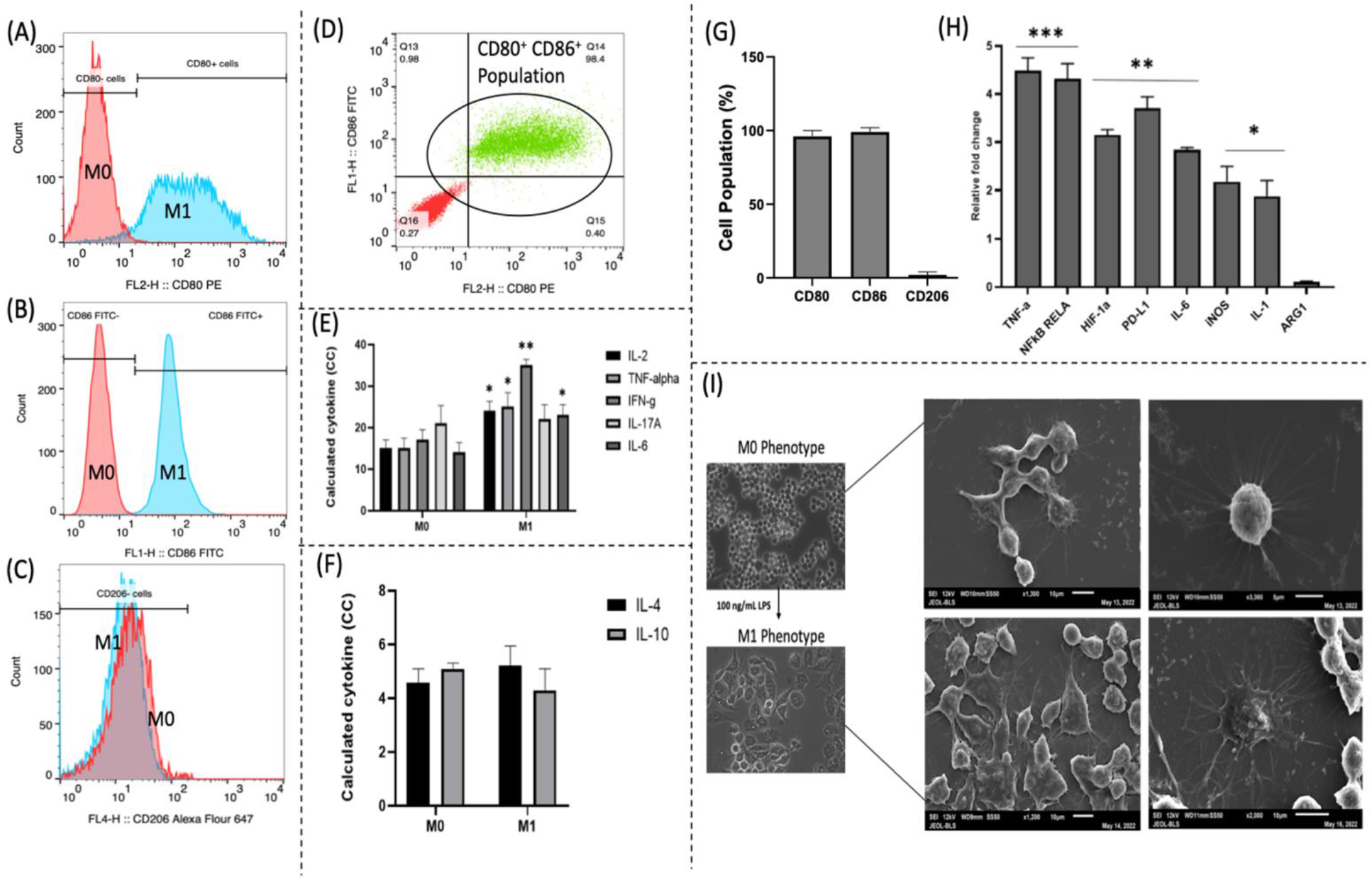
Pictorial and graphical representation of successful induction of M0 to M1 phenotype in RAW264.7 cells. (A) M1 macrophages exhibit higher CD80 expression, (B) M1 macrophages exhibit higher CD86 expression, (C) M1 macrophages express no changes in CD206 levels, (D) Dot plot depicting successful induction of CD80+ and CD86+ M1 subset, (E) Upregulation in proinflammatory cytokines by M1 macrophages, (F) No changes in anti-inflammatory cytokines by M1 macrophages, (G) RAW264.7 expressing each surface markers after LPS treatment, (H) Upregulation in expression levels of M1-like phenotypic markers, (I) Morphological changes observed in SEM after M1 to M2 induction. Mean ± SD, * p < 0.05, ** p < 0.01, *** p < 0.001, **** p < 0.0001, respectively.

This transition is crucial, as M1 macrophages are involved in inflammation initiation, phagocytosis, and antimicrobial defense. Increased levels of IL-2, IFN-γ, TNF-α, IL-17A, and IL-6 in M1 macrophages, compared to M0, were observed through cytokine bead array and confirmed by its gene expression by RT-PCR studies (Figure 11 E to H). These findings underscore the successful polarization of macrophages towards an M1-like phenotype, shedding light on the critical role M1 macrophages play in immune responses and their potential contribution to inflammation, phagocytosis, and antimicrobial defense. Figure 11 (I) illustrates morphological changes post-LPS treatment, showcasing amoeboid characteristics and delicate cytoplasmic extensions—hallmarks of M1 polarization associated with an activated and pro-inflammatory state. M1 macrophages play a pivotal role in initiating immune responses by secreting pro-inflammatory cytokines, chemokines, and reactive oxygen species to combat pathogens.

LPS leads to an increase in ROS production, however, with treatment with MTX + RELA siRNA FOL-liposome ranging from (25 nM to 100 nM), there was a significant decrease in ROS levels, indicating towards anti-inflammatory condition (Figure 12A). CD206, also known as the mannose receptor, signifies the M2 macrophage phenotype and is involved in glycoprotein endocytosis, immune regulation, and tissue repair. The observed decline in CD80 and CD86 expression indicates downregulated M1 markers, while increased CD206 expression confirms M2 marker upregulation, affirming the shift from an M1 to an M2 macrophage phenotype, at the lowest concentration, 25 nM of MTX + RELA siRNA FOL-liposome (Figure 12 B). This transition is vital for inflammation resolution, tissue repair, and immune regulation, ensuring proper tissue homeostasis and effective immune responses. Manipulating macrophage polarization holds therapeutic potential for various inflammatory and immune-related disorders. The heightened IL-2 levels in M1 macrophages signify increased production and secretion of cytokine, pivotal for T cell activation and immune response promotion. IFN-γ, predominantly from T cells and natural killer cells, underscores M1 macrophages’ activation and pro-inflammatory engagement.

**Figure 12.**
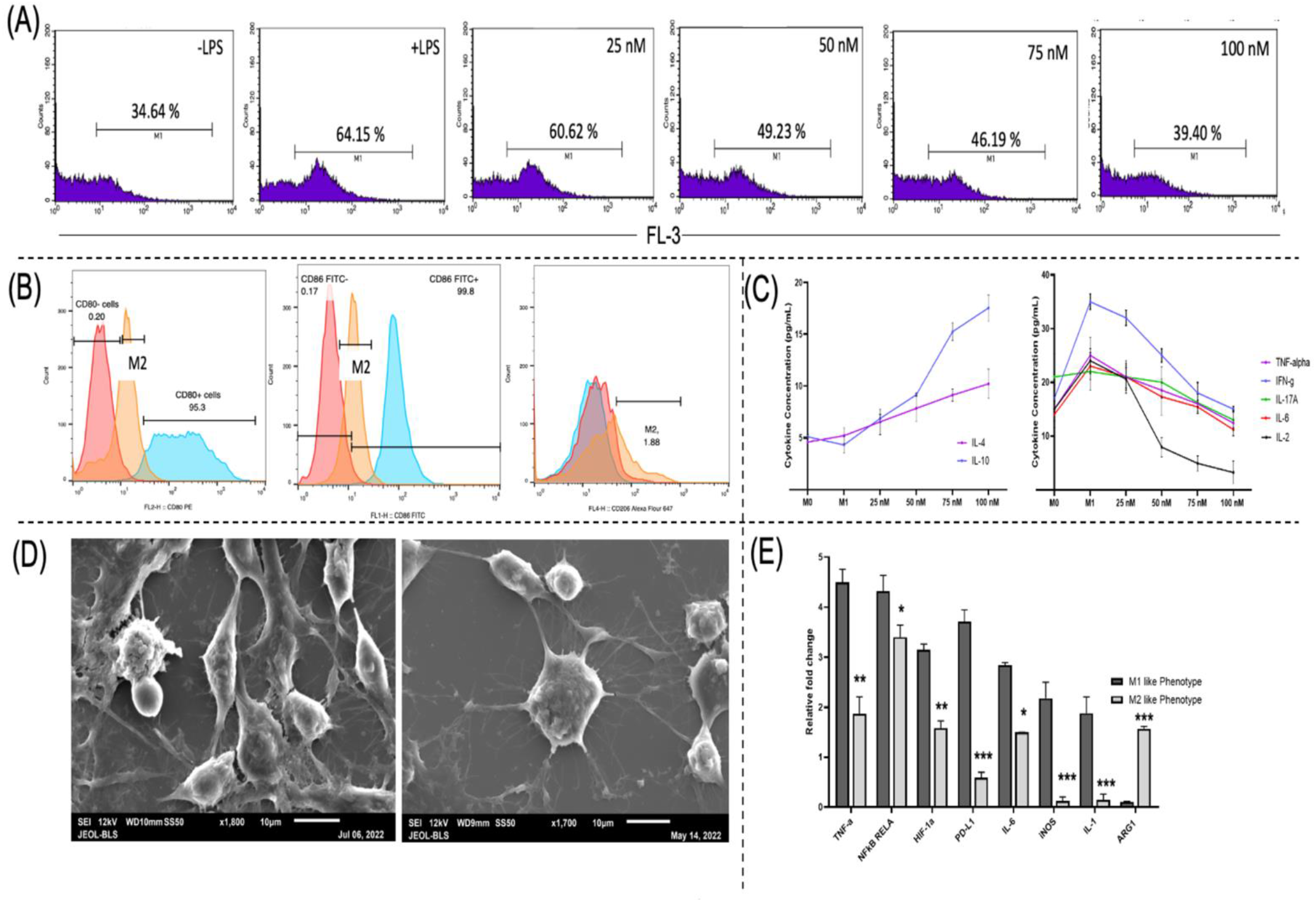
Pictorial and graphical representation of M1 to M2 polarization after the treatment of combinatorial folate liposome. (A) Concentration-dependent decrease in ROS, (B) Yellow peak depicts an M2-like population, and there is a decrease in CD80, and CD86 levels and a very slight increase in CD206 levels. (C) Concentration-dependent decrease in pro-inflammatory cytokines and increase in anti-inflammatory cytokines, (D) Dual shape of M2-like phenotype, ameboid and conical phenotypes, (E) Expression levels of M2-associated phenotype increases. Mean ± SD, * p < 0.05, ** p < 0.01, *** p < 0.001, **** p < 0.0001, respectively.

Elevated TNF-α in M1 macrophages indicates their role in fostering inflammation and immune activation, while heightened IL-17A suggests their contribution to inflammatory responses. IL-6’s increased levels in M1 macrophages imply participation in pro-inflammatory pathways. The levels of pro-inflammatory cytokines like TNF-α, IFN-γ, IL-17A, IL-6, and IL-2 decrease with the increasing treatment of MTX + RELA siRNA FOL-liposome, the anti-inflammatory cytokines like IL-4 and IL-10 increase due to MTX + RELA siRNA FOL-liposome. Notably, ARG-1, characteristic of M2 macrophages, was downregulated, highlighting the contrasting roles of M1 and M2 in immune regulation and tissue repair (Figures C and E). ARG-1, associated with tissue remodeling and wound healing, emphasizes M2 macrophages’ role in maintaining homeostasis (Figure 12 E). The balance between M1 and M2 macrophages is critical for effective immune responses and tissue health, with imbalances potentially contributing to disease pathology. Dual shape of M2-like phenotype, ameboid and conical were observed in macrophages after the treatment of MTX + RELA siRNA FOL-liposome (Figure 12 D).

Treatment-induced upregulation of secretory IL-4 and IL-10, along with decreased pro-inflammatory cytokines and increased ARG-1, signifies the M1 to M2 shift. IL-4 and IL-10, associated with the M2 phenotype, promote anti-inflammatory responses and tissue repair. Decreased pro-inflammatory cytokines align with the M1 to M2 transition, while ARG-1 upregulation supports M2 macrophages’ crucial role in tissue repair and immunoregulation.

### 2.6 *In vivo* studies in Collagen-induced Arthritic (CIA) Rats

The untreated control animals experienced a linear increase in growth, which is the natural growth pattern in the absence of any treatments or interventions. Animals receiving single treatments began to recover weight after 21 days (Figure 13 A). This indicates that the single treatments had some effect on alleviating the arthritic symptoms and allowing the animals to regain weight. However, the recovery was slower compared to the REL-A siRNA + MTX FOL-Liposome treated groups. Rats treated with a combination of REL-A siRNA and MTX loaded in FOL-Liposomes showed the most significant improvement. These treated groups recovered weight faster than the single treatment groups, indicating a more effective therapeutic response. The animals with CIA-induced arthritis experienced an increase in paw volume and arthritic scores due to edema formation, which is a common characteristic of inflammation (Figure B). Oedema refers to the accumulation of fluid in tissues, leading to swelling and pain in the joints. The combination liposomal treatment (REL-A siRNA + MTX FOL-Liposome) led to a decrease in edema formation, which contributed to a reduction in paw volume and an improvement in arthritic scores (Figure 13 C and D). This suggests that the combination treatment effectively mitigated the inflammatory response and reduced edema in the arthritic rats.

**Figure 13.**
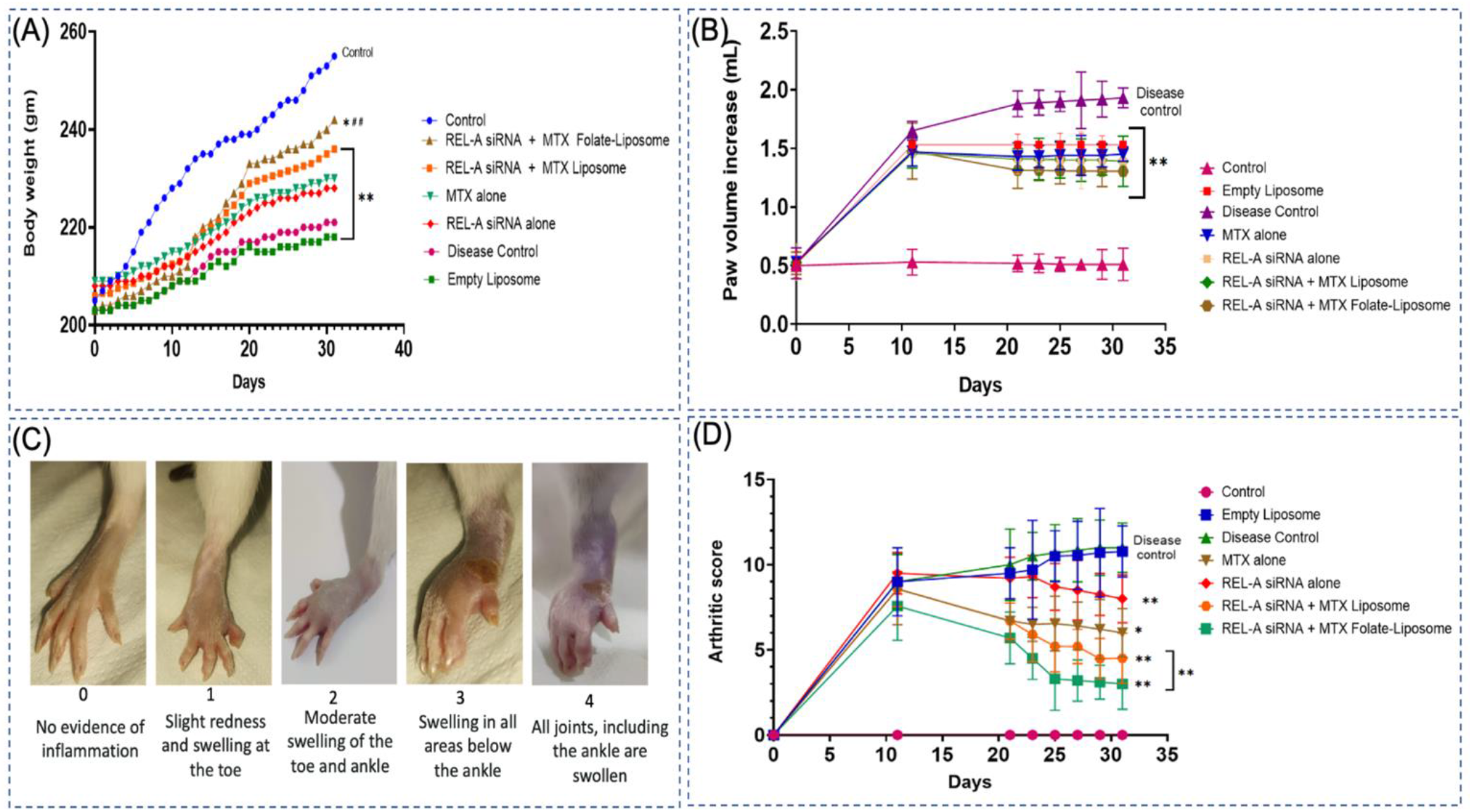
Pictorial and graphical representation of body weight fluctuation in all the 7 groups, % increase in paw volume, and Arthritic scores over 30 days. Groups= 7, n=3, Mean ± SD, * p < 0.05, ** p < 0.01, *** p < 0.001, **** p < 0.0001, with respect to placebo control. # p < 0.05, with respect to the disease control.

**Figure 14.**
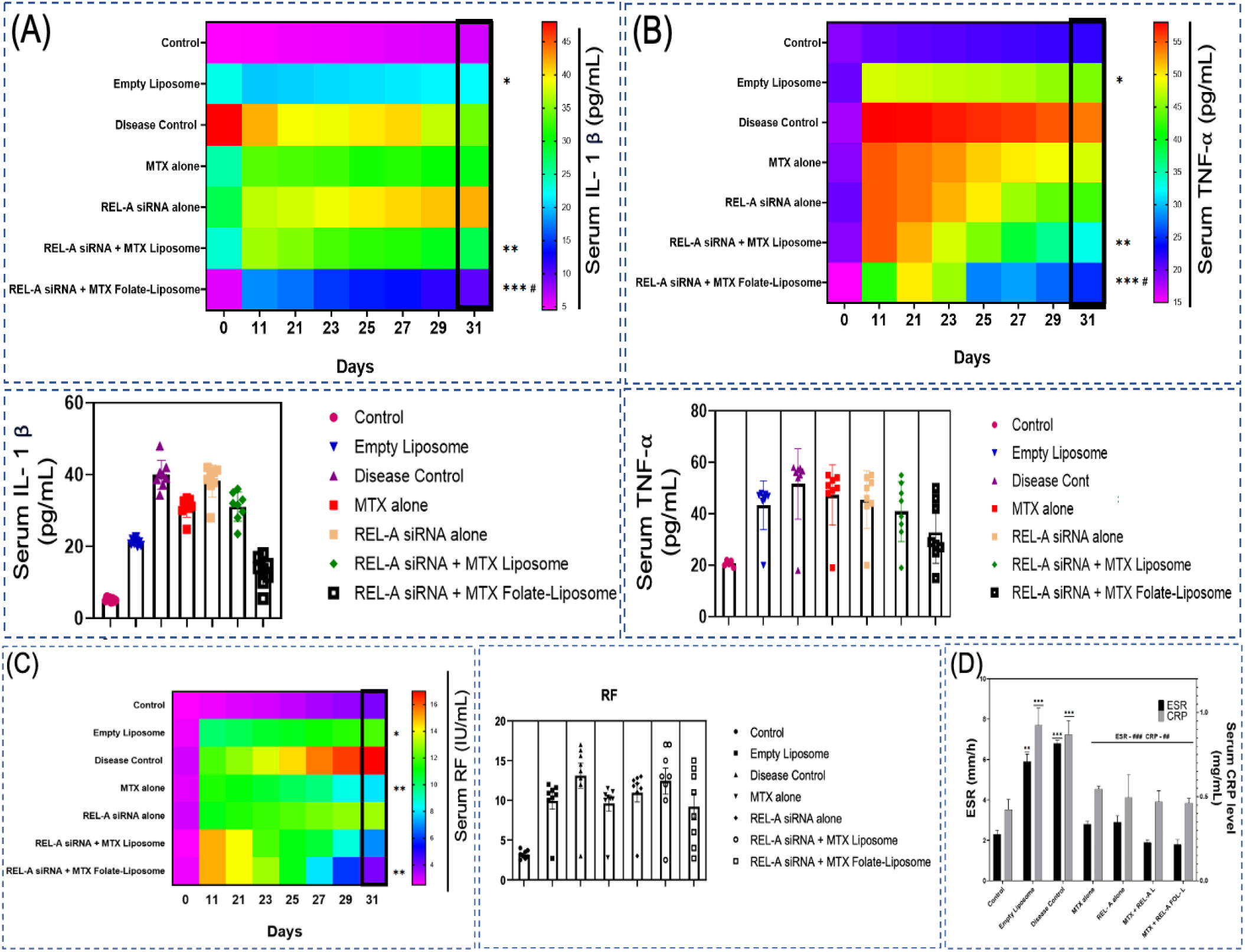
Graphical representation of elevated Serum TNF-α, Rheumatoid Factor (RF), Erythrocyte Sedimentation Rate (ESR), and C-reactive Protein (CRP) levels in CIA disease control group and subsequent reduction in these levels post treatments. Groups= 7, n=3, Mean ± SD, * p < 0.05, ** p < 0.01, *** p < 0.001, **** p < 0.0001, with respect to placebo control. # p < 0.05, with respect to disease control.

**Figure 15.**
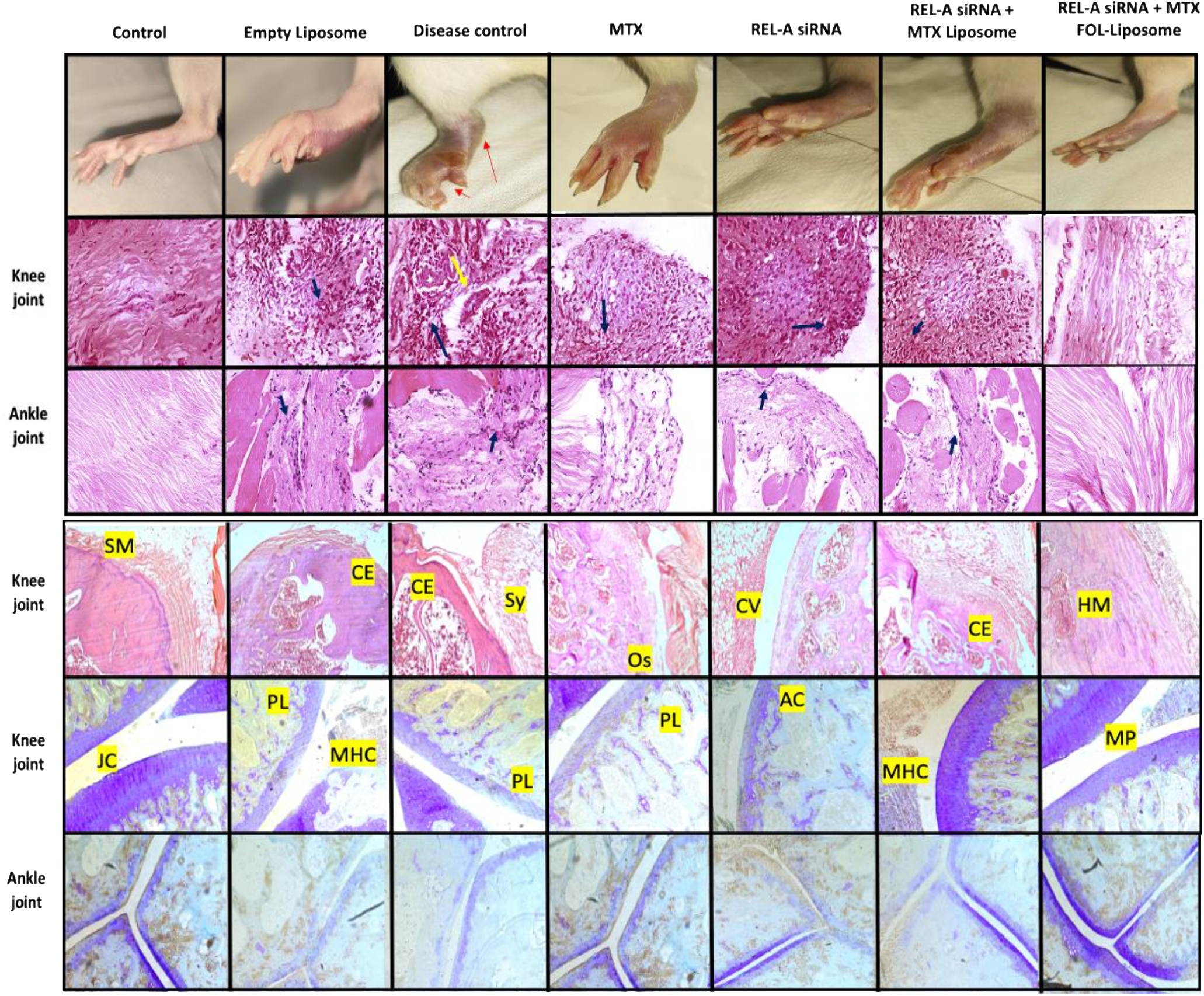
Pictorial representation of H&E and toluidine blue staining of ankle and knee joints in all 7 seven groups. Red arrows= excessive swelling, yellow arrow= Hypertrophy of Synoviocytes, Blue arrows= Lymphoplasmacytic Infiltration, SM= Synovial membrane, CE= Cartilage erosion, Sy= Synovitis, Os= Osteolysis, CV= Cellular vacuolization, HM= Hypercellular Marrow, JC= Joint cavity, PL= Proteoglycan loss, MHC= Meniscal Histopathological Changes, AC= Atypical Chondrocytes, and MP= Meniscal Proteoglycan Deficiency. Groups= 7 and n=3.

#### 2.6.1 Biochemical Parameters in Collagen-Induced Arthritic Rats

Tumor Necrosis Factor-alpha (TNF-α) is a pivotal pro-inflammatory cytokine implicated in RA pathogenesis, reflecting increased inflammation and joint damage. Rheumatoid Factor (RF) serves as an autoantibody, indicating disease severity and joint damage. Erythrocyte Sedimentation Rate (ESR) and C-reactive protein (CRP) are nonspecific inflammation markers, elevated in rheumatoid arthritis (Figure A to D). Monitoring TNF-α, RF, ESR, and CRP aids healthcare professionals in assessing disease activity, inflammation, and treatment response in rheumatoid arthritis patients.

#### 2.6.2 Histopathological Parameters in Collagen-Induced Arthritic Rats

The histopathological observations indicate marked signs of inflammation in the synovial membrane, which is a characteristic feature of inflammatory joint diseases, such as RA. Hypertrophy of Synoviocytes (yellow arrow): Synoviocytes are cells that line the synovial membrane and play a role in maintaining synovial fluid and joint function. The presence of hypertrophied synoviocytes, indicated by the yellow arrow, suggests abnormal enlargement and increased activity of these cells. This hypertrophy is a common response to inflammation and is a hallmark of inflammatory joint diseases. Diffuse Lymphoplasmacytic Infiltration (blue arrows): Lymphocytes and plasma cells are types of immune cells that play a significant role in the immune response. The presence of diffuse lymphoplasmacytic infiltration, indicated by the blue arrows, indicates the accumulation of these immune cells in the synovial membrane. Such infiltration is characteristic of inflammatory joint conditions, where immune cells migrate to the inflamed site and contribute to the inflammatory process.

Histopathological analysis of the disease control rat reveals active inflammation, characterized by hypertrophied synoviocytes and diffuse lymphoplasmacytic infiltration in the synovial membrane—features consistent with inflammatory joint diseases like rheumatoid arthritis. Examining synovial membrane histology is vital for diagnosing and understanding inflammation and tissue damage in joint diseases. This assessment aids healthcare professionals in gauging disease severity, monitoring progression, and evaluating therapeutic intervention efficacy. The use of toluidine blue, a metachromatic dye, highlights its utility in staining acidic components like proteoglycans, crucial in cartilage and connective tissues. The observed replenishment of proteoglycans suggests positive effects on joint tissues, potentially in the ankle joint, by MTX and RELA siRNA FOL-liposome treatment. This observation may suggest that the treatment is promoting cartilage health and helping to restore the joint’s natural protective and cushioning properties.

#### 2.6.3 X-ray Radiographic Analysis of Transtibial Joints of Hind Legs and Mobility Studies in CIA Rats

CIA rats in the disease control group experienced progressive destruction of the joints in their foot, including the joint between the tibia and tarsals. This joint destruction is a characteristic feature of inflammatory joint diseases like rheumatoid arthritis. X-ray images of the affected joints showed a reduction in the joint space, indicating cartilage loss and joint destruction. Cartilage acts as a cushion between bones in a joint, and its loss leads to the bones rubbing against each other, resulting in pain and reduced joint function. X-ray images also revealed soft tissue swelling as increased soft tissue density surrounding the joints. Soft tissue swelling is a common feature of joint inflammation and indicates the presence of fluid and cellular infiltration in the affected area. Difficulty in CIA rats in the disease control group experienced difficulty in mobility due to increased joint stiffness. Joint stiffness is a common symptom of inflammatory joint diseases and can severely impact an animal’s ability to move freely. Primary lesions refer to the initial site of damage or inflammation, while secondary lesions are additional lesions that develop as a consequence of the primary lesion. In all the groups that received CIA induction, there was an increase in both primary and secondary lesions. This indicates a widespread and progressive inflammatory response affecting various joints and tissues (Figure 16).

**Figure 16.**
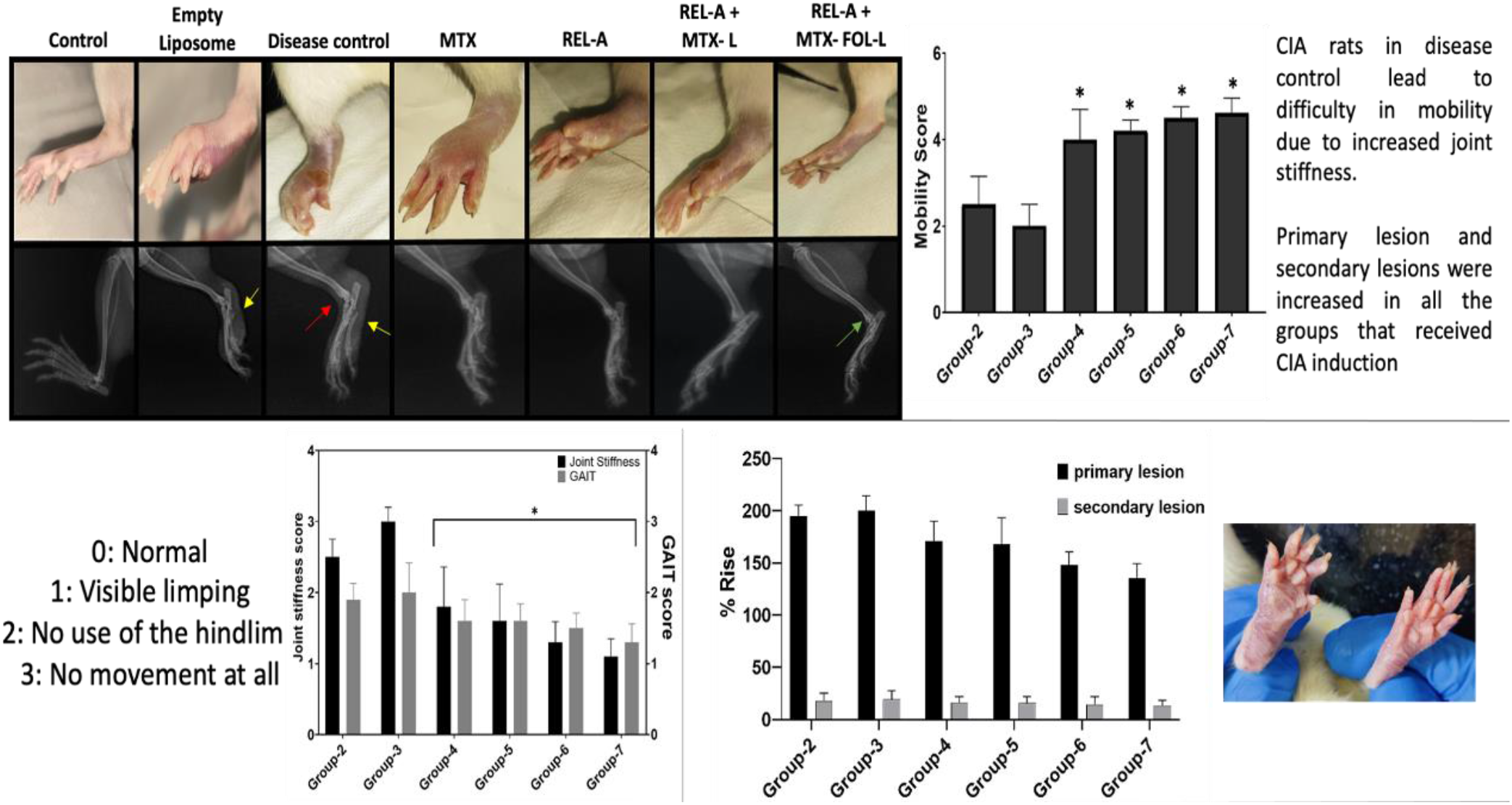
Radiographic analysis of transtibial joints of hind legs and mobility studies like GAIT score along with primary/secondary lesions. Yellow arrows suggest soft tissue swelling, the red arrow suggests a gap in the ankle joint and the green arrow suggests a reversal of the gap in the ankle joint. Groups= 7, n=3, Mean ± SD, * p < 0.05, ** p < 0.01, *** p < 0.001, **** p < 0.0001, with respect to placebo control.

**Figure 17.**
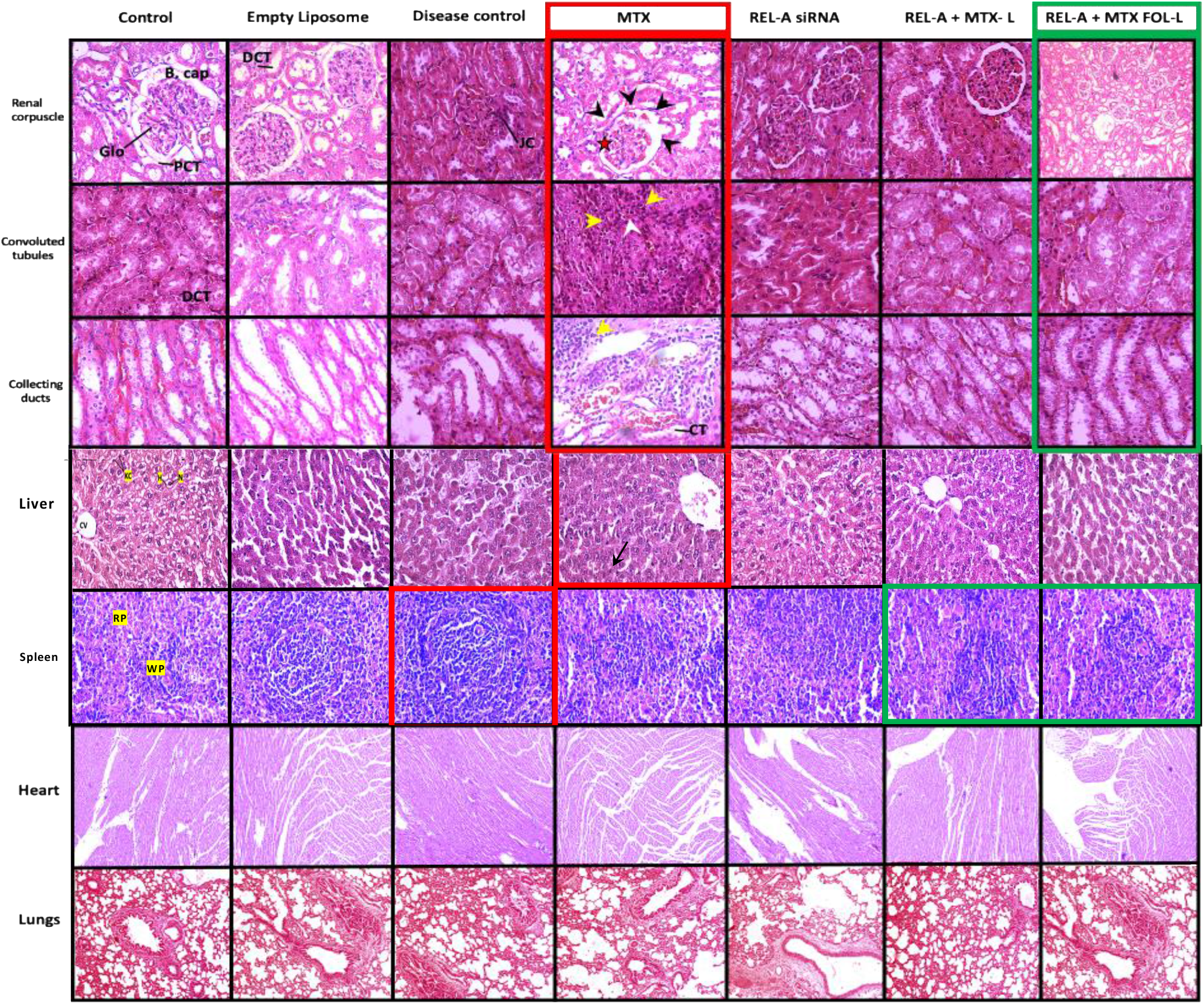
Pictorial representation of H&E stain assessing toxic effects and safety aspects in all the vital organs observed in all the 7 cohorts.

**Figure 18.**
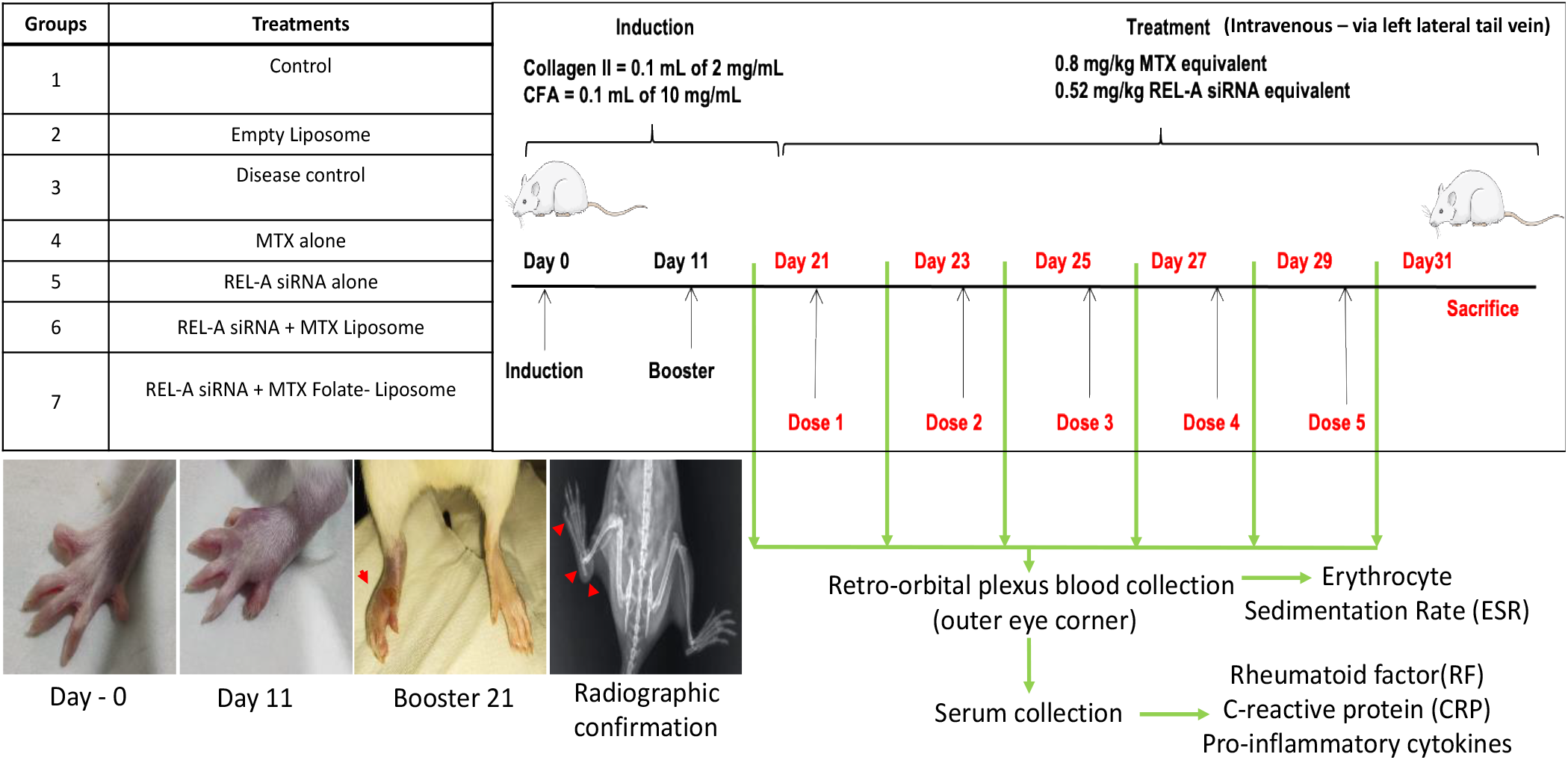
Animal grouping, Induction of CIA, and Treatment regimen. CFA-Complete Freund’s Adjuvant.

#### 2.6.3 Toxicological effects and safety assessment in each group

The MTX group in rats exhibited nephrotoxicity, which refers to the toxic effects of MTX on the kidneys. The researchers observed severe damage to the renal tissue, particularly in the renal cortex, but this damage was significantly reduced in group 7, suggesting a protective effect of the treatment in this group. In the MTX group, the researchers observed severe damage to the renal tissue, which is the kidney’s structural component. The renal cortex, the outer layer of the kidney, was particularly affected. The glomeruli are the filtration units of the kidneys, and their shrinkage suggests impaired kidney function. The dilatation of Bowman’s capsule, which surrounds the glomeruli, further indicates abnormal changes in the kidney’s filtration system. Inflammatory cell infiltration, including lymphocytes and plasma cells, was observed in the renal tissue. This infiltration is a sign of inflammation in the kidneys. Tubular degeneration suggests damage to the renal tubules, which are essential for reabsorption and secretion processes in the kidney. The presence of congestion in renal blood vessels indicates impaired blood flow in the kidneys, which can further contribute to kidney damage. The interstitial space of the kidney is the area between the renal tubules, and the infiltration of inflammatory cells (including lymphocytes and plasma cells) suggests ongoing inflammation in this area. The findings suggest that the MTX treatment led to significant nephrotoxicity in the rats, causing damage to the kidney’s structural components and impaired kidney function. However, in group 7, which likely received the combination treatment of REL-A siRNA and MTX FOL-Liposomes, these markers of nephrotoxicity were significantly reduced. This suggests that the combination treatment offered protection against the nephrotoxic effects of MTX, resulting in less severe kidney damage and inflammation.

Protecting against nephrotoxicity is crucial when using MTX for therapeutic purposes, as kidney damage can lead to serious complications. The observed reduction in nephrotoxicity markers in group 7 is promising and highlights the potential benefit of the combination treatment in mitigating the adverse effects of MTX on the kidneys. In the liver section, most of the hepatocytes (H) appeared to have a normal appearance, indicating that they were functioning normally and maintaining the liver’s normal structure and function. The nuclei (N) of hepatocytes were observed, suggesting that the cells were intact and had their nuclei, which is essential for their normal functioning. The central vein (CV), which is a blood vessel located in the centre of the liver lobule, was observed. It plays a role in carrying blood away from the liver and is an essential part of the liver’s vascular system. In the MTX group, a few pyknotic nuclei (black arrow) were observed. Pyknotic nuclei are condensed and shrunken nuclei, often seen in cells undergoing cell death or apoptosis. The presence of pyknotic nuclei in the MTX group suggests that MTX may have induced some cell death in the liver. In the spleen section, there was an observation of aggravated white pulp (WP) after RA induction. The spleen is an important organ of the immune system, and its white pulp contains clusters of immune cells, such as lymphocytes, involved in immune responses. The aggravation of the white pulp indicates an increased immune response, likely in response to RA induction.

Overall, the histopathological findings in the liver showed a normal appearance of most hepatocytes, but with a few pyknotic nuclei in the MTX group, suggesting some liver cell death. In the spleen, RA induction led to aggravated white pulp, indicating an increased immune response.

The above figure also describes the safety assessment of the formulated liposomes on vital organs like lungs and heart isolated from a healthy control group and the control + Liposome treated groups. The healthy control group, which did not receive any liposome treatment, served as the reference for normal histoarchitecture of the vital organs. The liposome treatment was administered to the control group to assess its potential effects on vital organs. The H&E staining of the control + Liposome treated groups showed no histoarchitectural changes in vital organs such as the heart, lungs, and liver. This indicates that the liposome treatment did not cause any apparent damage or disruption to the normal structure of these organs. The histology of the vital organs in the control + Liposome treated groups was found to be similar to that of the healthy control group. This suggests that the liposome treatment did not induce any noticeable differences in tissue morphology compared to the untreated healthy organs.

## 3. Conclusion

The study showcased the antiarthritic potential of Methotrexate (MTX) and RELA siRNA within FOL-Liposomes, employing extensive in vitro and in vivo analyses. Thorough liposome characterization revealed stability and high biocompatibility. FOL-Liposomes exhibited quasi-spherical morphology, sustained MTX release, and remarkable entrapment efficiency for MTX and RELA siRNA. In a rat model of inflammatory arthritis, RELA siRNA reduced synovial inflammation, while MTX and RELA siRNA in FOL-Liposomes indirectly inhibited cytokines, RF, and CRP, improving mobility. Targeting macrophages via folate receptors revealed a synergistic therapeutic impact, potentially shifting macrophage phenotype and preventing cartilage erosion. This nano-formulation holds promise for innovative inflammatory arthritis management, suppressing inflammation and preserving joint integrity.

## 4. Experimental section

### 4.1 Materials and Chemicals

2-distearoyl-sn-glycero-3-phosphoethanolamine-N-[meth-oxy(polyethyleneglycol)-2000-folate (DSPE-PEG2000-folate)], 1,2-distearoyl-sn-glycero-3-phosphoethanolamine-N-[methoxy(polyethyleneglycol)-2000] (DSPEPEG2000), 1,2-Distearoyl-sn-glycero-3-phosphatidylcholine (DSPC), and cholesterol were procured from Avanti Polar Lipids, India. Dimethyl sulfoxide (DMSO), LPS, 2, 7-dichlorofluorescein diacetate (DCF-DA), 3-[4, 5-dimethylthiazol-2-yl]-2, 5-diphenyl-tetrazolium bromide (MTT), Dulbecco’s Modified Eagle’s Medium (DMEM), Antibiotic Antimycotic Solution 100x Liquid, Fetal Bovine Serum (FBS), were obtained from Gibco™ -Thermo Fisher Scientific. The Cytokine Bead Array kit was procured from BD Biosciences. Primers for murine macrophage were purchased from Eurofins Scientific, India. Rhodamine dye, Fluorescein isothiocyanate (FITC), 4′,6-diamidino-2-phenylindole (DAPI), and Phalloidin-Atto 488 were obtained from Sigma-Aldrich. PE Hamster Anti-Mouse CD80 (BD PharmingenTM), FITC Rat Anti-Mouse CD86 (BD PharmingenTM), Alexa Fluor-647 Rat Anti-Mouse CD206 (BD PharmingenTM) and Glutaraldehyde solution, 25% w/w was obtained from HiMedia. Freund’s complete adjuvant and Collagen Type II were obtained from Sigma-Aldrich.

### 4.2 Synthesis and Characterization of MTX + RELA siRNA FOL-liposomes

DSPC, Cholesterol, DSPE-PEG, and DSPE-PEG-FOL were used to synthesize liposomes in the molar ratio as derived in the 3 factorial Box Behnken design (DSPC/CHOL = 4:1, Total Lipids/MTX = 10, and DSPE-PEG-FOL = 0.06). The lipids were dissolved in chloroform and further rotated under pressure till a thin film was formed, using a rotary vacuum evaporator model EV11, at 65°C [21]. Thin film formation was followed by 1 h hydration with 2 mL of phosphate buffer saline (PBS) containing calcium phosphate nanoparticle and RELA siRNA and loading of 100 µL of 1mg/mL MTX, at 55-60 °C. The suspension was extruded 11 times through a 100 nm filter using an Avanti Polar extruder. The prepared MTX-Liposomes were stored at 4 °C and were characterized. The Zetasizer (Malvern Instruments) was used to determine the particle size and polydispersity index of formulations at 25°C using DLS. Liposomes were tested for encapsulation efficiency, particle hydrodynamic size, and polydispersity index after 30 days of storage at 4°C for this study.

### 4.3 Characterization of liposomes

The hydrodynamic size, zeta potential, and polydispersity index of the liposome formulations were evaluated using the Zetasizer (Malvern Instruments Ltd. Malvern, UK), through Dynamic light scattering at 25 °C. The sample was freshly prepared before every read, by adding 10 microlitres of liposomes in 1 mL PBS. The stability of liposome formulations was monitored consistently for 30 days, and the liposomes were found to be stable at 4 °C.

### 4.4 Transmission Electron Microscopy (TEM)

The morphology of the liposomes was observed using a transmission electron microscope (JEOLJEM1400) at a 120/100 kV Voltage. Sample preparation was done using a negative stain method, where a drop of liposome formulation was stained with Phosphotungstic acid (PTA), on a TEM grid, and observed after overnight incubation.

### 4.5 Atomic force microscopy (AFM)

Freshly cleaved mica surfaces were used as substrates for AFM imaging. A small aliquot (typically 20-50 µL) of liposome formulation was gently deposited onto the mica surface. Samples were allowed to adsorb for 30-60 minutes in a humid environment to prevent drying artifacts. AFM imaging was performed using a high-resolution atomic force microscope (Bruker Nano Wizard Sense).

### 4.6 Fourier transform infrared (FTIR)

The FTIR analysis of the samples was done at room temperature, using the PerkinElmer spectrum. The samples were prepared by diluting several times, and lyophilization using KBr. The wavelength range of the scans was from 4000 to 400 cm-1, and the results were normalized for better analysis.

### 4.7 Confocal Imaging

Confocal microscopy imaging was performed using a confocal laser scanning microscope (e.g., Leica SP8). A suitable objective lens with 40X magnification was selected for imaging. Fluorescent labelling of liposomes was performed using Rhodamine, while Phalloidin and DAPI were used as cytoplasmic and nuclear stains, respectively. Z-stack images were captured to obtain three-dimensional information about the sample. Images were acquired as optical sections and later reconstructed into three-dimensional representations.

### 4.8 Determination of drug entrapment efficiency and kinetic release

Liposomes were stored inside the dialysis membrane and this assembly was then immersed in a beaker containing PBS with a pH of 6.5 while being continuously stirred. After a 12-hour incubation period, the PBS solution was collected to determine the entrapment efficiency. Subsequently, the PBS was replaced with fresh PBS (pH 6.5), and the setup was maintained for an additional 48 hours to collect hourly samples (1hr to 8hr, 12hr, 24hr, and 48hr) for drug release analysis. High-Performance Liquid Chromatography (HPLC) using a Shimadzu prominence-i instrument and a spectrophotometer was employed to quantify the drug content in the collected solutions. The amount of drug released within the initial 24 hours was utilized to calculate the Entrapment Efficiency (%EE), while the kinetic drug release profile was established.

### 4.9 MTT Assay

The cytotoxicity of the FOL-Liposomes, MTX-Liposomes, and MTX-FOL-liposomes was carried out on RAW264.7 cells. MTT Assay evaluated the percentage Cell proliferation of the final formulation. 7 × 10^3^/well M0 cells were seeded in 96-well culture plates and incubated overnight at 37 °C. Further, the Cells were stimulated with LPS (100 ng/mL) for 18 h, leading to M1 phenotype macrophages. After achieving the M1 phase, various concentration treatments of liposome formulations were given and incubated for 24 and 48 hours. After that cells were incubated for 3 h with serum-deprived media and 5mg/mL MTT. 0.5% MTT was used for each well. Further MTT containing medium was replaced with 100 µL DMSO and the absorbance was taken using a spectrophotometer (Synergy HT, Bio-Tek) with 5 min shaking at 570 nm wavelength.

### 4.10 Cellular Surface Markers Analysis

Cells with 50×10^3^ density were seeded in a 6-well culture plate. After overnight incubation cells were treated with 100 ng/mL LPS for 18 hours to achieve the M1 phenotype. Treatments were given and incubated for 24 hours. Further cells were incubated for 3 h with a cocktail of CD206, CD86, and CD80 antibodies. Cell fixation and mounting were performed using PFA (Paraformaldehyde) and Moviol respectively. The slides were analyzed under confocal microscopy.

### 4.11 Gene Expression Studies using qRT-PCR

Total RNA from untreated M0 cells, LPS (100 ng/ml) treated M1 cells, and MTX + RELA siRNA treated M2 cells was extracted using the Favourogen RNA Isolation kit, according to the manufacturer’s protocol. The purity and concentration of the isolated RNA were estimated using a SYNERGY-HT multiwell plate reader (Synergy HT, Bio-Tek, USA) using the absorbance ratio 260/280. The isolated RNA was reverse-transcribed into cDNA by the Maxima first-strand cDNA synthesis kit, according to the manufacturer’s protocol. The real-time PCR of the synthesized cDNA was accomplished using the SYBR Green Master Mix (applied biosystems); along with the sense and antisense primers for all the inflammatory marker genes, with GAPDH as a housekeeping gene. PCR was performed in the Eppendorf thermocycler for 40 cycles. Gene expression was quantified by the ΔΔCT method using Quant Studio 5.

### 4.12 CIA Induction and Treatment

The experiments were performed at LM College of Pharmacy (LMCP), Ahmedabad. The Female Wistar rats, weighing around 180−200 g, were kept in standard conditions under guidelines provided by CPCSEA in an animal house of LMCP. Initially, before treatment, they were acclimatized for 6 days. The disease induction was started on the seventh day (Refer to supplementary information file for detailed protocol and methodology for *in vivo* experiments).

### 4.13 Statistical analysis

The experiments were carried out three times. The data were presented as the mean standard deviation for outcomes that were continuous. One-way and two-way analysis of variance (ANOVA) using GraphPad Prism was used to find statistically significant differences (P<0.05) between the three groups (version 5; La Jolla, CA, USA).

## Supporting Information

Supporting Information is available from the Wiley Online Library or from the author.

## Notes

The authors declare no competing financial interest.

## Author Contribution

SN: Investigation, Experimentation, Methodology, Writing - Original draft. DB: Experimentation, Writing - review and editing AK: Conceptualization, Writing - review and editing, Supervision, Resources, and Funding acquisition.

## Animal ethical approval

All the experiments were performed following the guidelines provided by the Institutional Animal Ethics Committee. The approval number for this project is LMCP/IAEC/22/0044.

## Acknowledgement

We would like to acknowledge the Gujarat State Biotechnology Mission (GSBTM) (GSBTM/JD(R&D)/663/2023-24/02003699) and The Gujarat Institute for Chemical Technology (GICT) (AU/DBLS/GICT-nanomaterials/2016-17/01) for funding this work. SN acknowledges Ahmedabad University for providing PhD fellowship.

## Abbreviations

AFM: Atomic force microscopy
TEM: Transmission electron microscopy
FTIR: Fourier transform infrared spectroscopy
TNF-*α*: Tumor necrosis factor-*α*
NF-*κ*B: Nuclear factor kappa B
MTX: Methotrexate
siRNA: short interfering RNA.

## Notes

### Competing Interest Statement

The authors have declared no competing interest.

## References

1. Zyrianova, Y., Rheumatoid arthritis-etiology, consequences and co-morbidities 2012, 9, 807–11.

2. Gersh, B. J.; Sliwa, K.; Mayosi, B. M.; Yusuf, S., European heart journal 2010, 31 (6), 642–648.

3. Kłak, A.; Raciborski, F.; Samel-Kowalik, P., Reumatologia/Rheumatology 2016, 54 (2), 73–78.

4. Malm, K.; Bergman, S.; Andersson, M. L.; Bremander, A.; Larsson, I., SAGE open medicine 2017, 5, 2050312117713647.

5. Drosos, A. A.; Pelechas, E.; Voulgari, P. V., Clinical Rheumatology 2020, 39 (4), 1363–1368.

6. Bevaart, L.; Vervoordeldonk, M. J.; Tak, P. P., Arthritis Rheum 2010, 62 (8), 2192–2205.

7. Nalwa, H. S.; Prasad, P.; Ganguly, N. K.; Chaturvedi, V.; Mittal, S. A., Translational Medicine Communications 2023, 8 (1), 1–9.

8. Mustafa, S. H.; Ahmad, T.; Balouch, M.; Iqbal, F.; Durrani, T., Cureus 2022, 14 (7).

9. Friedman, B.; Cronstein, B., Joint bone spine 2019, 86 (3), 301–307.

10. Conforti, A.; Di Cola, I.; Pavlych, V.; Ruscitti, P.; Berardicurti, O.; Ursini, F.; Giacomelli, R.; Cipriani, P., Autoimmunity reviews 2021, 20 (2), 102735.

11. Yang, X.; Chang, Y.; Wei, W., Cell proliferation 2020, 53 (7), e12854.

12. Nasra, S.; Bhatia, D.; Kumar, A., Nanoscale Advances 2022, 4 (17), 3479–3494.

13. Wang, N.; Liang, H.; Zen, K., Frontiers in immunology 2014, 5, 614.

14. Fujiwara, N.; Kobayashi, K., Current Drug Targets-Inflammation & Allergy 2005, 4 (3), 281–286.

15. Nasra, S.; Shah, T.; Bhatt, M.; Chaudhari, R.; Bhatia, D.; Kumar, A., ACS Applied Bio Materials 2023, 6 (7), 2886–2897.

16. Komano, Y.; Yagi, N.; Onoue, I.; Kaneko, K.; Miyasaka, N.; Nanki, T., Journal of Pharmacology and Experimental Therapeutics 2012, 340 (1), 109–113.

17. Bitounis, D.; Fanciullino, R.; Iliadis, A.; Ciccolini, J., International Scholarly Research Notices 2012, 2012.

18. Mortazavi, S. A., 2014.

19. Sedky, N. K.; Braoudaki, M.; Mahdy, N. K.; Amin, K.; Fawzy, I. M.; Efthimiadou, E. K.; Youness, R. A.; Fahmy, S. A., Nanoscale Advances 2023, 5 (19), 5399–5413.

20. Ji, Y.; Yang, X.; Ji, Z.; Zhu, L.; Ma, N.; Chen, D.; Jia, X.; Tang, J.; Cao, Y., ACS omega 2020, 5 (15), 8572–8578.

21. Andra, V. V. S. N. L.; Pammi, S.; Bhatraju, L. V. K. P.; Ruddaraju, L. K., BioNanoScience 2022, 12 (1), 274–291.

