## Supplementary Information for "Synergistic Modulation of Macrophages by Methotrexate and RELA siRNA Folate-Liposome: A Precision Therapy to Prevent Joint Degradation in Collagen-Induced Arthritic Rats"

*Corresponding Author:

Ashutosh Kumar,

Associate Professor

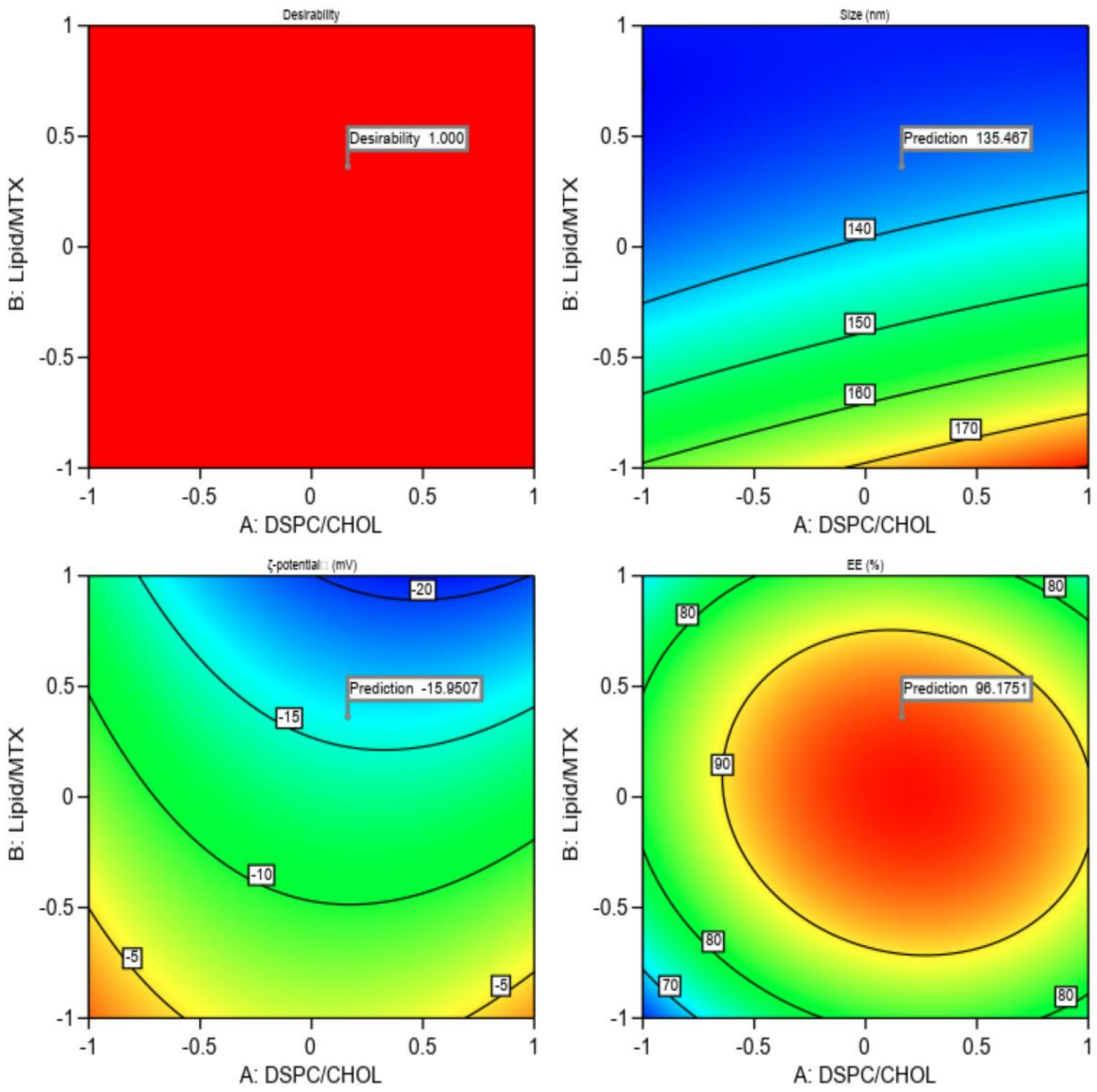


**Figure S1.** Desirability score 1 suggests that liposomes have a hydrodynamic size of 135.5 nm, zeta potential of around -15.95, and Entrapment Efficiency of 96.175 % are most preferable, which aligns with formulation 5, mentioned in Table S1.

| Run | DSPC/CHOL | Lipid/MTX | DSPE-PEG-FOL | Response 1 %EE | Response 2 Size | Response 3 Z-potential |
|---|---|---|---|---|---|---|
| 1 | 1 | 0 | 0 | 93.25 | 130.8 | -10.9 |
| 2 | 0 | 0 | 0 | 89.6 | 140.2 | -12.7 |
| 3 | 0 | 0 | 0 | 98.39 | 140.8 | -12.9 |
| 4 | 0 | 1 | 0 | 82.3 | 133.1 | -21.22 |
| 5 | 0 | 0 | 1 | 98.39 | 140.8 | -12.9 |
| 6 | 0 | 0 | 0 | 89.6 | 140.2 | -12.7 |
| 7 | 1 | 0 | 0 | 98.072 | 147 | -11.87 |
| 8 | 0 | 0 | 0 | 98.39 | 140.8 | -12.9 |
| 9 | 0 | -1 | 0 | 79 | 169 | -6.38 |
| 10 | 1 | -1 | 1 | 80.4 | 181.3 | -3.97 |
| 11 | -1 | 1 | 1 | 74 | 130 | -9.96 |
| 12 | -1 | 0 | 0 | 88.8 | 134 | -8.6 |
| 13 | 1 | 1 | -1 | 82.8 | 143.8 | -20 |
| 14 | -1 | -1 | -1 | 62.02 | 148.2 | -0.256 |
| 15 | 0 | 0 | -1 | 98.2 | 120.4 | -10.2 |

**Table S1.** 15 runs for 3 independent variables and 3 responses.

Arthritis evaluation encompassed the assessment of various parameters:

1. Body Weight: Body weight was recorded weekly from day 0 to day 31.

2. Paw Volume: The plethysmometer was employed to measure the paw volume daily from the onset of disease in all animal groups.

3. Primary and Secondary Lesions:

    a. *Primary Lesion:* Edema formation in the injected hind paw peaking 3–5 days after the phlogistic agent injection. A percent increase in the injected left paw over control on day 31 was calculated.
    b. *Secondary Lesions:* Immunologically mediated changes characterized by inflammation in non-injected sites. The evaluation involved calculating the percent increase in the non-injected right paw over control on day 31 [1].

4. Arthritic Scores: The severity of arthritis in each limb was assessed using the following scoring system:
    a. 0—No evidence of erythema or swelling
    b. 1—Mild swelling or erythema largely present in joints
    c. 2—Mild swelling or erythema extending from the ankle to tarsals
    d. 3—Erythema and moderate swelling extending from the ankle to metatarsal joints
    e. 4—Erythema and severe swelling encompassing the ankle, foot, digits, or ankyloses of the limb. Scores were recorded daily [2].

5. Joint Stiffness: On day 31, joint stiffness was scored by manipulating ankle bending and extension within the limits of the range of motion. Scores were assigned based on restrictions: full range of movement in both bending and extension (0), restriction in bending or extension (1), or restriction in both (2) [3].

6. Gait Test: The use of the untreated (right) paw was scored on day 31, considering creeping behavior (2), passive use to support the body (1), or active use (0). Scores from two observers were combined for each rat and totaled for group scores [3].

7. Mobility Test: On day 31, mobility scoring included [3]:
    a. Walks normally (6)
    b. Walks are protective toward the ipsilateral hind paw (5)
    c. Walks are protective toward the ipsilateral hind paw, touching only the toe (4)
    d. Walks being protective toward both hind paws, touching fully or only the toe (3)
    e. Crawls only use the fore paws (1)
    f. Does not move (0).
8. Biochemical Parameters: On the 31st day, rats were anesthetized, and blood samples were collected for:
    a. Erythrocyte sedimentation rate (ESR) using the Wintrobe method
    b. C-reactive protein (CRP) and rheumatoid factor (RF) measurement by turbidimetric method using commercial kits (Beacon Diagnostics Ltd, India).
    c. Serum TNF-a and IL-1 β measurement using a rat ELISA kit (Weldon Biotech Pvt. Ltd., India) as per the kit manual.

9. X-ray Visualization. X-ray analysis was performed to assess the arthritis severity in the paws and joints of rats. All the rats were anesthetized using mild anesthesia and kept on a radiosensitive plate 110 cm distant from the X-ray source. Images were taken by a Portable Veterinary X-ray machine with a generator capacity of 100 mA and high frequency.

10. Hematoxylin and eosin staining: Rats were subjected to isoflurane anesthesia and euthanized on the 31st day of the experiment. Subsequently, their ankle joints were extracted and preserved in 4% formalin. Following decalcification in 5% formic acid, the joints underwent paraffin embedding. Sections of 5µm thickness were cut and subjected to hematoxylin and eosin staining, then examined using a bright field microscope [4].

11. Toluidine blue staining: 5µm thick joint sections were deparaffinized and gradually rehydrated in decreasing ethanol concentrations, followed by water. The sections were

then stained with 0.4% toluidine blue and counterstained with 0.001% fast green (FCF). Afterward, the sections underwent dehydration using increasing ethanol concentrations and xylene. Finally, the sections were mounted with DPX and observed under a bright field microscope at 10X magnification [5].

References:

(1) Philippe, L.; Gegout-Pottie, P.; Guingamp, C.; Bordji, K.; Terlain, B.; Netter, P.; Gillet, P. Relations between functional, inflammatory, and degenerative parameters during adjuvant arthritis in rats. *American Journal of Physiology-Regulatory, Integrative and Comparative Physiology* **1997**, *273* (4), R1550-R1556.
(2) Jia, Q.; Wang, T.; Wang, X.; Xu, H.; Liu, Y.; Wang, Y.; Shi, Q.; Liang, Q. Astragalin suppresses inflammatory responses and bone destruction in mice with collagen-induced arthritis and in human fibroblast-like synoviocytes. *Frontiers in pharmacology* **2019**, *10*, 94.
(3) Nagakura, Y.; Okada, M.; Kohara, A.; Kiso, T.; Toya, T.; Iwai, A.; Wanibuchi, F.; Yamaguchi, T. Allodynia and hyperalgesia in adjuvant-induced arthritic rats: time course of progression and efficacy of analgesics. *Journal of Pharmacology and Experimental Therapeutics* **2003**, *306* (2), 490-497.
(4) Ahmad, A.; Ansari, M. M.; Mishra, R. K.; Kumar, A.; Vyawahare, A.; Verma, R. K.; Raza, S. S.; Khan, R. Enteric-coated gelatin nanoparticles mediated oral delivery of 5-aminosalicylic acid alleviates severity of DSS-induced ulcerative colitis. *Materials Science and Engineering: C* **2021**, *119*, 111582.
(5) Zhang, Q.; Dehaini, D.; Zhang, Y.; Zhou, J.; Chen, X.; Zhang, L.; Fang, R. H.; Gao, W.; Zhang, L. Neutrophil membrane-coated nanoparticles inhibit synovial inflammation and alleviate joint damage in inflammatory arthritis. *Nature nanotechnology* **2018**, *13* (12), 1182-1190.
